# Magnesium modulates phospholipid metabolism to promote bacterial phenotypic resistance to antibiotics

**DOI:** 10.1101/2024.06.24.600343

**Authors:** Hui Li, Jun Yang, Su-fang Kuang, Huan-zhe Fu, Hui-ying Lin, Bo Peng

**Affiliations:** State Key Laboratory of Bio-Control, School of Life Sciences, Guangdong Province Key Laboratory for Pharmaceutical Functional Genes, Southern Marine Science and Engineering Guangdong Laboratory (Zhuhai), Sun Yat-sen University, Higher Education Mega Center, Guangzhou 510006, People’s Republic of China; Laboratory for Marine Biology and Biotechnology, Marine Fisheries Science and Food Production Processes, Qingdao Marine Science and Technology Center, Qingdao 266071, China

**Keywords:** magnesium, phenotypic resistance, quinolones, phospholipid metabolism, membrane fluidity

## Abstract

Non-inheritable antibiotic or phenotypic resistance ensures bacterial survival during antibiotic treatment. However, exogenous factors promoting phenotypic resistance are poorly defined. Here, we demonstrate that *Vibrio alginolyticus* are recalcitrant to killing by a broad spectrum of antibiotics under high magnesium. Functional metabolomics demonstrated that magnesium modulates fatty acid biosynthesis by increasing saturated fatty acid biosynthesis while decreasing unsaturated fatty acid production. Exogenous supplementation of unsaturated and saturated fatty acids increased and decreased bacterial susceptibility to antibiotics, respectively, confirming the role of fatty acids in antibiotic resistance. Functional lipidomics revealed that glycerophospholipid metabolism is the major metabolic pathway remodeled by magnesium, where phosphatidylethanolamine (PE) biosynthesis is reduced and phosphatidylglycerol (PG) production is increased. This process alters membrane composition, increasing membrane polarization, and decreasing permeability and fluidity, thereby reducing antibiotic uptake by *V. alginolyticus*. These findings suggest the presence of a previously unrecognized metabolic mechanism by which bacteria escape antibiotic killing through the use of an environmental factor.

## Introduction

Non-inheritable antibiotic or phenotypic resistance represents a serious challenge for treating bacterial infections. Phenotypic resistance does not involve genetic mutations Phenotypic resistance does not involve genetic mutations and is transient, allowing bacteria to resume normal growth. Biofilm and bacterial persisters are two phenotypic resistance types that have been extensively studied (Brandis *et al*., 2023; Corona & Martinez, 2013). Biofilms have complex structures, containing elements that impede antibiotic diffusion, sequestering and inhibiting their activity (Ciofu *et al*., 2022). Biofilm-forming bacteria and persisters also have distinct metabolic states that significantly reduce their antibiotic susceptibility (Yan & Bassler, 2019). These two types of phenotypic resistance share the common feature in their retarded or even cease of growth in the presence of antibiotics (Corona & Martinez, 2013). However, specific factors that promote phenotypic resistance and allow bacteria to proliferate in the presence of antibiotics remain poorly defined.

Metal ions have a diverse impact on the chemical, physical, and physiological processes of antibiotic resistance (Booth *et al*, 2011; Lu *et al*, 2020; Poole, 2017). This includes genetic elements that confer resistance to metals and antibiotics (Poole, 2017) and metal cations that directly hinder (or enhance) the activity of specific antibiotic drugs (Zhang *et al*., 2014). The metabolic environment can also impact the sensitivity of bacteria to antibiotics (Jiang *et al*., 2023; Lee & Collins, 2012; Peng *et al*., 2015; Zhang *et al*., 2020; Zhao *et al*., 2021). Light metal ions, such as magnesium, sodium, and potassium, can behave as cofactors for different enzymes (Du *et al*., 2016) and influence drug efficacy. Heavy metal ions, including Cu^2+^ and Zn^2+^, confer resistance to antibiotics (Yazdankhah *et al*., 2014; Zhang *et al*., 2018). Recent reports suggest that sodium negatively regulates redox states to promote the antibiotic resistance of *Vibrio alginolyticus* (Yang *et al*., 2018), while actively growing *Bacillus subtilis* cope with ribosome-targeting antibiotics by modulating ion flux (Lee *et al*, 2019). In Gram-negative bacteria, by contrast, zinc enhances antibiotic efficacy by potentiating carbapenem, fluoroquinolone, and β-lactam-mediated killing (Isaei *et al*., 2016; Zhang *et al*., 2014). Magnesium influences bacterial structure, cell motility, enzyme function, cell signaling, and pathogenesis (Wang *et al*., 2019). This mineral also modulates microbiota to harvest energy from the diet (Garcia-Legorreta *et al*., 2020), allowing *Bacillus subtilis* to cope with ribosome-targeting antibiotics by modulating ion flux (Lee *et al*., 2019). However, the role of magnesium in promoting phenotypic resistance is less well understood.

Vibrios inhabit seawater, estuaries, bays, and coastal waters, regions full of metal ions such as magnesium (Kumarage *et al*., 2022). Magnesium is the second most dissolved element in seawater after sodium. At a salinity of 3.5% seawater, the magnesium concentration is about 54 mM (Potis, 1968), and in deep seawater, can be as high as 2,500 mM (Wang *et al*., 2024). *Vibrio parahaemolyticus* and *V. alginilyticus* are two representative Vibrio pathogens that infect humans and aquatic animals, resulting in illness and economic loss, respectively (Grimes, 2020). (Fluoro)quinolones such as balofloxacin are used to treat Vibrio infection, however, resistance has emerged due to overuse (Suyamud *et al*., 2024). Indeed, (fluoro)quinolones are one of China’s two primary residual chemicals associated with aquaculture (Liu *et al*., 2017). Vibrio can develop quinolone resistance through mutations in the DNA gyrase gene or through plasmid-mediated mechanisms (Dutta *et al*., 2021). Thus, the use of *V. parahaemolyticus* and *V. alginilyticus* as bacterial representatives, and balofloxacin as a quinolone-based antibacterial representative, can help to define novel magnesium-dependent phenotypic resistance mechanisms of pathogenic Vibrio species.

The current study evaluated whether magnesium induces phenotypic resistance in Vibrio species and defined the molecular/genetic basis for this resistance. Genetic approaches, GC-MS analysis of metabolite and membrane remodeling upon antibiotic exposure, membrane physiology, and extensive antimicrobial susceptibility testing were used for the evaluations.

## Results

### Mg^2+^ promotes phenotypic resistance to antibiotics

Marine environments and agriculture, where antibiotics are commonly used, are rich in magnesium. To investigate whether this mineral impacts antibiotic activity, the minimal inhibitory concentration (MIC) of *V. alginolyticus* ATCC33787 and *V. parahaemolyticus* VP01, which we referred as ATCC33787 and VP01 afterwards, isolated from marine aquaculture, to balofloxacin (BLFX) in Luria-Bertani medium (LB medium) plus 3% NaCl as LBS medium and “artificial seawater” (ASWT) medium that included the major ion species in marine water (Wilson, 1975) (LB medium plus 210 mM NaCl, 35 mM Mg_2_SO_4_, 7 mM KCl, and 7 mM CaCl_2_) were assessed (**Suppl. Table 1**). The MICs of ATCC33787 to BLFX were 54 and 1 μg/mL in ASWT and LBS medium, respectively. Similarly, the MICs to VP01 were 25 and 3 μg/mL in ASWT and LBS medium, respectively (**Fig 1A**). The role of exogenous NaCl in antibiotic resistance was investigated previously (Yang *et al*., 2018). The current study found that the MIC for BLFX was the same in LB and M9 minimal medium (M9 medium) plus 7 mM KCl or 7 mM CaCl_2_ (**Suppl. Fig 1**). However, the MIC for BLFX was higher in ASWT medium supplemented with Mg_2_SO_4_ or MgCl_2_ than in LB medium (**Fig 1B**). And Mg_2_SO_4_ or MgCl_2_ had no difference on MIC, suggesting it is Mg^2+^ not other ions contribute to the MIC change. The MIC for BLFX increased at higher concentrations of MgCl_2_ in ASWT medium. Specifically, adding 50 mM or 200 mM MgCl_2_ increased the MIC for BLFX by 200- or 1600-fold, respectively (**Fig 1C**). At balofloxacin doses of 1.56, 3.125, 6.25, and 12.5 µg, the zone of inhibition decreased with increasing MgCl_2_ (**Fig 1D**). Exogenous MgCl_2_ also increased the MICs for other quinolone (*e.g.* nalidixic acid, levofloxacin, ciprofloxacin, ofloxacin, and moxifloxacin) (**Fig 1E**) and non-quinolone antibiotics including antibacterial peptides (colistin), macrolides (roxithromycin), tetracyclines (oxytetraycline), β-lactams (ceftriaxone, ceftazidime), and aminoglycosides (amikacin, kanamycin, and gentamicin) (**Fig 1F**). Notably, magnesium had a reduced effect on ceftriaxone and gentamicin than other antibiotics. Importantly, exogenous MgCl_2_ also increased MICs of clinic isolates, carbapenem-resistant *Escherichia coli*, carbapenem-resistant *Klebsiella pneumoniae,* carbapenem-resistant *Pseudomonas aeruginosa* and carbapenem-resistant *Acinetobacter baumannii* to balofloxacin (**Fig 1G**). These findings indicate that Mg^2+^ promotes phenotypic resistance.

**Figure 1.**
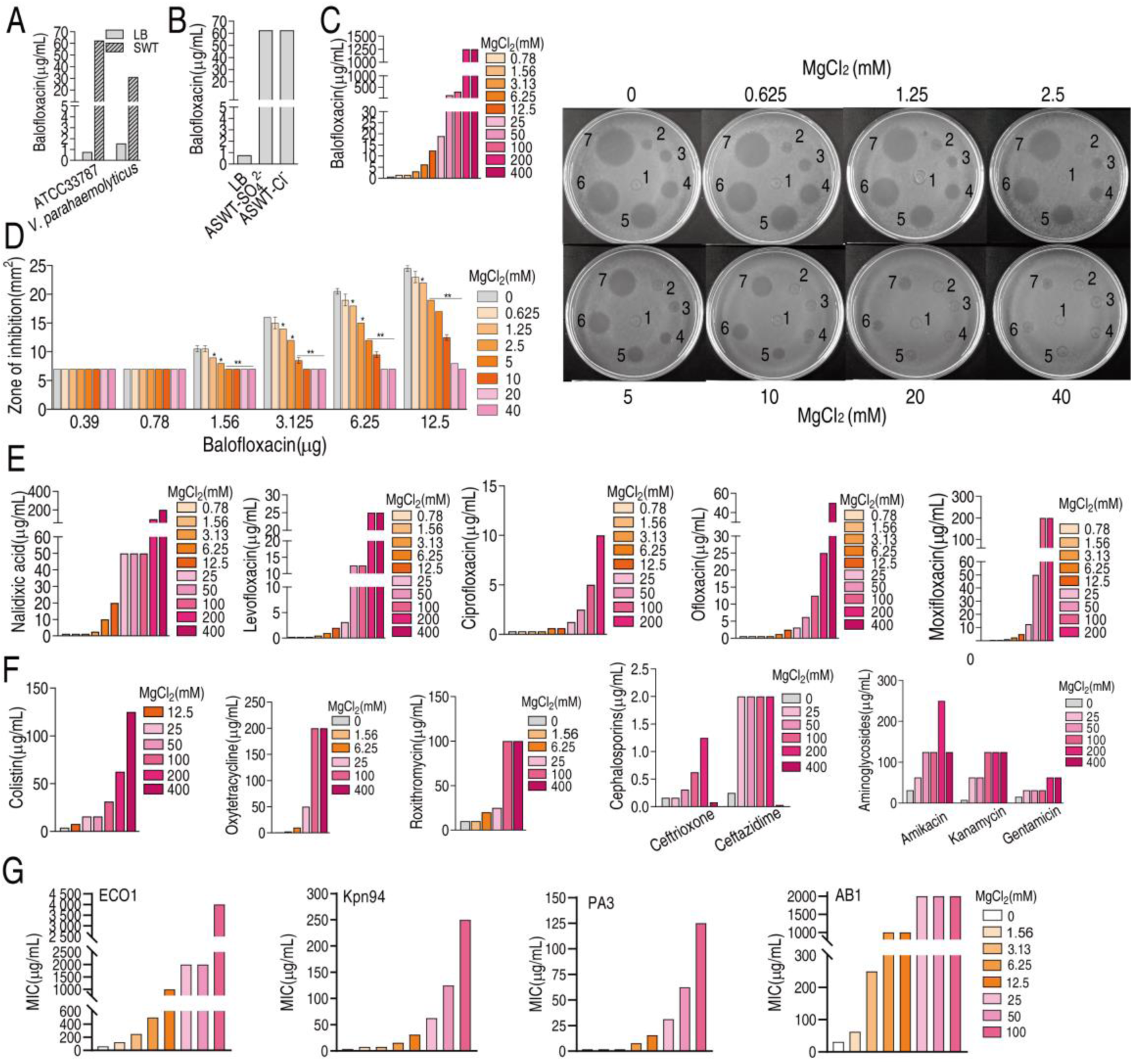
Magnesium promotes bacterial resistance to antibiotics. **A.** MIC of ATCC33787 and *V. parahaemolyticus* to BLFX in ASWT or LB medium as determined by the microtitre-dilution method. **B.** MIC of ATCC33787 to BLFX in ASWT with additional MgSO_4_ or MgCl_2_ as determined by the microtitre-dilution-method. **C.** MIC of ATCC33787 to BLFX in ASWT at the indicated concentrations of MgCl_2_ as determined by the microtitre-dilution method. **D.** MIC of ATCC33787 to BLFX at the indicated concentrations of BLFX and MgCl_2_ as determined by the Oxford cup test. The numbers 1, 2, 3, 4, 5, 6, and 7 represent 0, 0.39, 0.78, 1.56, 3.125, 6.25, and 12.5 µg BLFX, respectively. **E.** MIC of ATCC33787 to other quinolones in ASWT at the indicated concentrations of MgCl_2_ as determined by the microtitre-dilution method. **F.** MIC of ATCC 33787 to other classes of antibiotics in ASWT at the indicated concentrations of MgCl_2_ as determined by the microtitre-dilution method. **G.** MIC of carbapenem-resistant *Escherichia coli*, carbapenem-resistant *Klebsiella pneumoniae,* carbapenem-resistant *Pseudomonas aeruginosa,* and carbapenem-resistant *Acinetobacter baumannii* isolates to BFLX at the indicated concentrations of MgCl_2_.

### MgCl_2_ affects bacterial metabolism

To investigate how magnesium regulates phenotypic resistance, other effects of magnesium, including its ability to quench balofloxacin activity and induce efflux pump expression and LPS biosynthesis, were excluded (**Suppl. Text; Suppl. Fig 2**). While Mg^2+^ is a co-factor for enzymes (Garfinkel and Garfinkel, 1985), whether this mineral can promote antibiotic resistance independent of this function remains unknown.

To better understand how magnesium affects bacterial metabolism, *V. alginolyticus* was cultured in M9 medium in the presence of various amounts of MgCl_2_ (0, 0.78, 3.125, 12.5, 50, or 200 mM) and the levels of 54 metabolites were assessed by GC-MS. Five biological and two technical replicates were evaluated for each treatment (**Suppl. Fig 3**). The levels of 41 metabolites were differential (p <0.05) and are presented as Z-values (**Fig 2A; Suppl. Fig 4)**. Orthogonal partial least square discriminant analysis (OPLS-DA) was conducted that separated the six treatments into three groups: Group 1 included 0 mM, 0.78 mM, and 3.125 mM MgCl_2_; Group 2 was a singlet of 12.5 mM; Group 3 included 50 mM and 200 mM MgCl_2_. Component t[1] distinguishes Group 1 from Groups 2 and 3; component t[2] distinguishes Group 3 from Groups 1 and 2 (**Fig 2B**). Discriminating variables are shown as S-plot, where cut-off values are ≥ 0.05 absolute value of covariance p and 0.5 for correlation p(corr). Six biomarkers/metabolites were selected from component t[1] and p[1]. More specifically, the abundance of cadaverine, urea, palmitic acid, aminoethanol, and fumaric acid were increased, but pyroglutamic acid and glutamic acid were decreased (**Fig 2C; Suppl. Fig 5A**). Pathway enrichment analysis suggests that twelve pathways are involved. Notably, one of the pathways is biosynthesis of unsaturated fatty acids (terminology in the software. Actually, it is biosynthesis of fatty acids) (**Fig 2D**), where the abundance of palmitic acid and stearic acid were increased in an Mg^2+^ dose-dependent manner (**Fig 2E; Suppl. Fig 6**). Moreover, palmitic acid was a crucial biomarker (**Suppl. Fig 6**). The increase in fatty acid biosynthesis could be partially explained by an imbalanced pyruvate cycle/TCA cycle, in which fumarate levels increased at higher Mg^2+^ while succinate levels increased at lower Mg^2+^ (**Suppl. Fig 5B**). These findings indicated that glycolysis fluxes into fatty acid biosynthesis rather than the pyruvate cycle/TCA cycle. The relevance of fatty acids and BLFX was demonstrated by the observation that exogenous palmitic acid increased bacterial resistance to balofloxacin (**Fig 2F**). These results suggest that fatty acid metabolism may be critical to magnesium-based phenotypic resistance.

**Figure 2.**
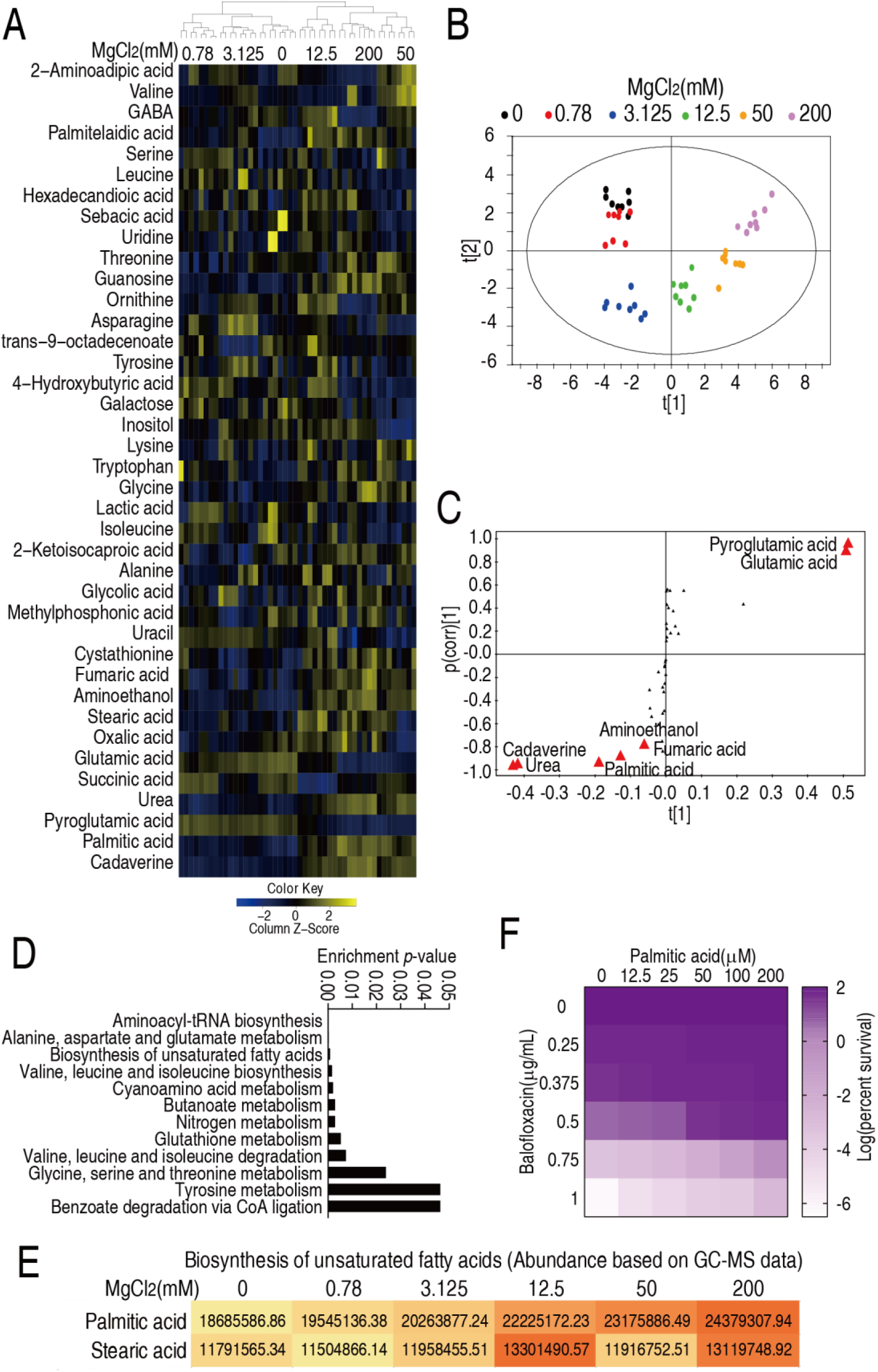
Mg^2+^-induced metabolomic change. **A.** Differential metabolomes in the absence or presence of the indicated concentrations of MgCl_2._ The yellow and blue colors indicate an increase or decrease in metabolite levels relative to the median metabolite level, respectively (see color scale). Euclidean distance calculations were used to generate a heatmap that shows clustering of the biological and technical replicates of each treatment. **B.** OPLS-DA analysis of different MgCl_2_-induced metabolome concentrations. Each dot represents a technical replicate of samples in the plot. **C.** S-plot generated from OPLS-DA. Predictive component p [1] and correlation p(corr) [1] differentiate 0, 0.78, 3.125 mM MgCl_2_ from 12.5, 50, 200 mM MgCl_2_. Predictive component p[2] and correlation p(corr)[2] separate 0, 0.78, 50, 200 mM MgCl_2_ from 3.125, 12.5 mM MgCl_2_. The triangle represents metabolites in which candidate biomarkers are marked. **D.** Enriched pathways by differential abundances of metabolites. **E.** Areas of the peaks of palmitic acid and stearic acid generated by GC-MS analysis. **F.** Synergy analysis for BFLX with palmitic acid for *V. alginolyticus*. Synergy was performed by comparing the dose needed for 50% inhibition of the synergistic agents (white) and non-synergistic (i.e., additive) agents (purple).

### Mg^2+^ regulates fatty acid biosynthesis

Acetyl-CoA carboxylase (ACC) catalyzes the conversion of acetyl-CoA to malonyl-CoA, the first step of fatty acid synthesis. Fatty acid biosynthesis, which includes saturated and unsaturated forms, shares common biosynthetic pathways with acetyl-CoA to enoyl-ACP, and is sequentially mediated by *fabD* or *fabH*, *fabB*/*F*, *fabG,* and *fabA*/*Z*. The enzyme, enoyl-ACP, produces unsaturated fatty acids or is metabolized to acyl-ACP via *fabV* to produce saturated fatty acids. Unsaturated fatty acids produce saturated fatty acids via *tesA*/*B* and *yciF* (**Fig 3A**). qRT-PCR was used to quantify the expression of involved genes in bacteria treated with different concentrations of MgCl_2_. The expression of 18 genes increased following treatment with 50 and/or 200 mM MgCl_2_ (**Fig 3B**). The expression of *tesA*, *tesB*, and *yciA*, which convert unsaturated to saturated fatty acids, increased at 50 or/and 200 mM MgCl_2_ (**Fig 3C**). FabA and FabF play key roles in unsaturated fatty acid and fatty acid biosynthesis, respectively (Feng & Cronan, 2009; Lee *et al*., 2013). Western blot analysis showed that FabA levels declined and FabF levels increased at higher MgCl_2_ concentrations (**Fig 3D**). Oxazole-2-amine and triclosan inhibit the synthesis of fatty acids and decrease bacterial survival in the presence of balofloxacin (**Fig 3E and 3F**). These results indicate that Mg^2+^ inhibits the synthesis of unsaturated fatty acids and promotes the synthesis of saturated fatty acids.

**Figure 3.**
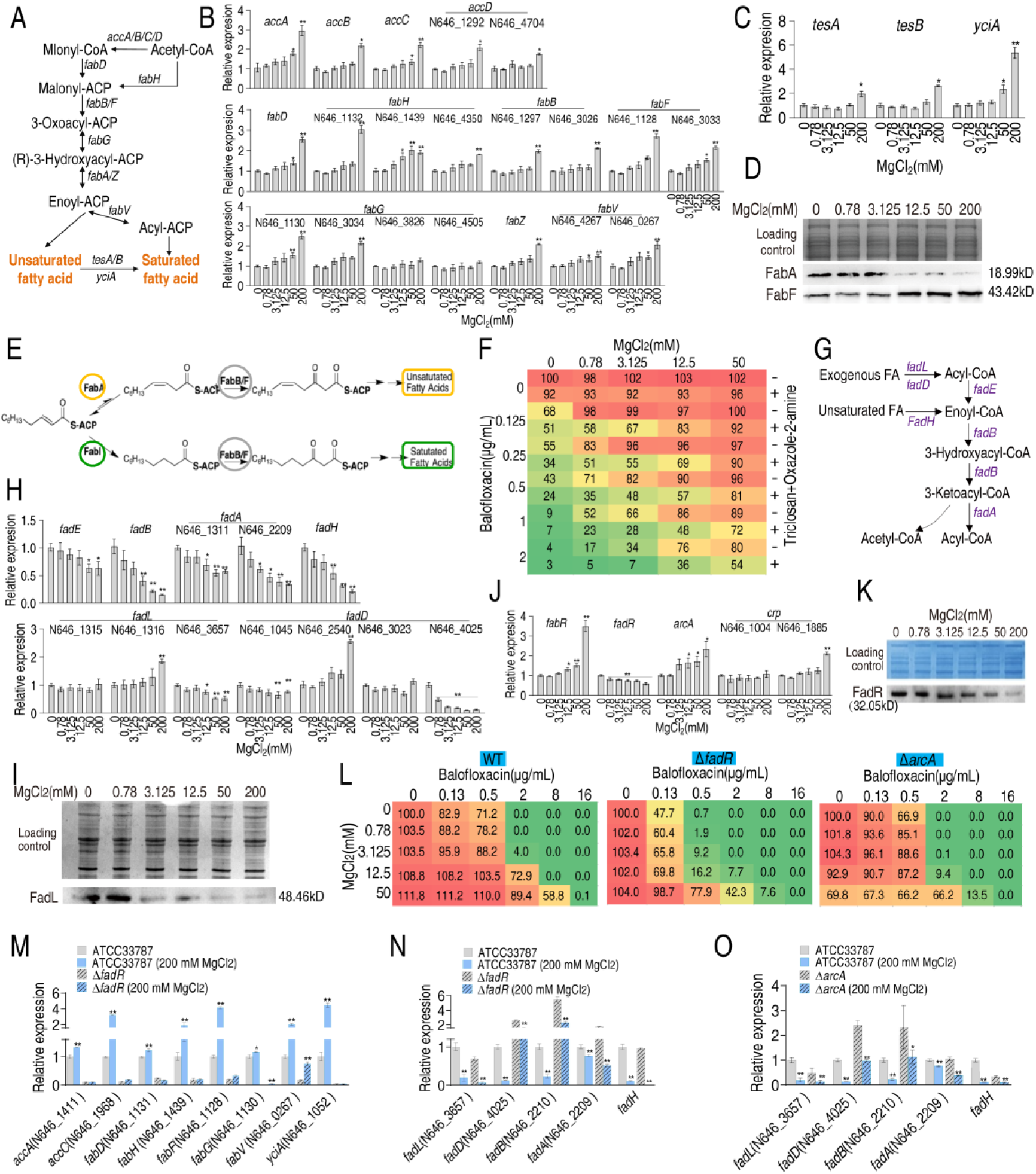
Mg^2+^ promotes fatty acid biosynthesis. **A.** Diagram of fatty acid biosynthesis. **B.** qRT-PCR for expression of fatty acid biosynthesis genes in the absence or presence of indicated concentrations of MgCl_2_. **C.** qRT-PCR for expression of genes involved in converting unsaturated fatty acids to saturated fatty acids in the absence or presence of indicated MgCl_2_ levels. **D.** Western blot for the abundance of proteins responsible for converting unsaturated fatty acids to saturated fatty acids in the absence or presence of indicated concentrations of MgCl_2_. **E.** Diagram of saturated and unsaturated acid biosynthesis. **F.** Synergy analysis of balofloxacin with triclosan + oxazole-2-amine for ATCC33787. The percentage of bacterial survival was quantified in the presence of indicated MgCl_2_ levels and/or balofloxacin plus triclosan and oxazole-2-amine and used to construct the isobologram. Synergy is represented using a color scale or an isobologram, which compares the dose needed for 50% inhibition of synergistic agents (blue) and non-synergistic (i.e., additive) agents (red). **G.** Diagram of fatty acid degradation. **H.** qRT-PCR for the expression of genes encoding fatty acid degradation in the absence or presence of the indicated concentrations of MgCl_2_. **I.** Western blot for the abundance of FadL in the absence or presence of the indicated concentrations of MgCl_2_. **J.** qRT-PCR for the expression of fatty acid biosynthesis regulatory genes in the absence or presence of the indicated concentrations of MgCl_2_. **K.** Western blot for the abundance of FadR in the absence or presence of the indicated concentrations of MgCl_2_. **L.** Synergy analysis for MgCl_2_ with BLFX for ATCC33787 (WT), Δ*fadR*, and Δ*arcA*. Synergy is represented using a color scale or an isobologram, which compares the dose needed for 50% inhibition of the synergistic agents (blue) and non-synergistic (i.e., additive) agents (red). **M, N,** and **O** qRT-PCR for expression of genes involved in fatty acid biosynthesis (**M**) and degradation **(N** and **O)** in ATCC33787, Δ*fadR* or/and Δ*arcA* **(O)** in the presence or absence of 200 mM MgCl_2_. Whole cell lysates resolved by SDS-PAGE gel was stained with Coomassie brilliant blue as loading control **(D)**, **(I) and (K)**.

In addition, we also quantified gene expression during fatty acid degradation to determine whether Mg^2+^ affects this process (**Fig 3G and 3H**). Interestingly, the expression of genes involved in unsaturated fatty acid degradation decreased in an MgCl_2_ dose-dependent manner (**Fig 3H**). Western blot analysis confirmed that FadL levels decreased with increasing MgCl_2_ (**Fig 3I**). FadH degrades unsaturated fatty acids and influences the ratio of unsaturated to saturated fatty acids. Together, these results suggest that MgCl_2_ inhibits fatty acid degradation.

FabR, FadR, ArcA, and cAMP/CRP are transcriptional factors involved in regulating fatty acid biosynthesis. FabR inhibits unsaturated fatty acid biosynthesis, FadR promotes fatty acid biosynthesis and inhibits degradation, and ArcA and cAMP/CRP inhibit and promote fatty acid degradation, respectively (Feng and Cronan, 2010; Fujita et al., 2007). The expression of *fabR* and *arcA* increased, while the expression of *fadR* decreased. The expression of N646_1885, which encodes CRP, increased only in the presence of 200 mM MgCl_2_ (**Fig 3J**). Western blot analysis confirmed that FadR levels declined with increasing MgCl_2_ concentration (**Fig 3K**). Viability of Δ*fadR* and Δ*arcA* was reduced in the presence of MgCl_2_, balofloxacin, or both in a dose-dependent manner (**Fig 3L**). Thus, FadR and ArcA may play roles in bacterial resistance to balofloxacin, presumably by a mechanism dependent on the abundance of saturated/unsaturated fatty acids.

To investigate whether Mg^2+^ functions through FadR and ArcA, expression of eight fatty acid biosynthetic genes (*accA, accC, fabD, fabH, fabR, fabG, fabV,* and *yciA*) and five fatty acid degradation genes (*fadL, fadD, fadB, fadA,* and *fadH*) were quantified in wildtype, Δ*fadR* or/and Δ*arcA* bacterial strains. As expected, fatty acid biosynthetic gene expression was reduced in Δ*fadR* cells (**Fig 3M**). While higher expression of *fadD, fadB,* and *fadA* (elevated *fadA* only for Δ*fadR* without 200 mM MgCl_2_) was detected in Δ*fadR* or Δ*acrA* strains, the expression of these genes was negatively regulated by 200 mM MgCl_2_ (**Fig 3N and 3O**). These results indicate that Mg^2+^ promotes fatty acid biosynthesis by positively regulating *fadR*, and inhibits the degradation of fatty acids through *fadR* and *arcA*.

### MgCl_2_ affects 16-carbon and 18-carbon fatty acid metabolism

To investigate the effect of Mg^2+^ on the biosynthesis of saturated and unsaturated fatty acids, LC-MS was used to quantify the levels of 16-carbon and 18-carbon fatty acids, the main precursors of lipid biosynthesis (Zhang and Rock, 2008b). The abundance of the saturated fatty acid, palmitic acid (C16:0), was higher while the abundance of five unsaturated fatty acids, palmitoleic acid (C16:1), linoelaidic aid (C18:2), linoleic acid (C18:2), alpha-linoleic acid (C18:3), and stearidonic acid (C18:4) was reduced with increasing MgCl_2_ concentration (**Fig 4A**). The levels of stearic acid (C16:0) and vaccenic acid (C18:1) remained unaffected. Total saturated fatty acid levels increased and total unsaturated fatty acid levels decreased with increasing Mg^2+^ (**Fig 4B**). These results suggest that Mg^2+^ upregulates saturated fatty acid biosynthesis but downregulates unsaturated fatty acid biosynthesis.

**Figure 4.**
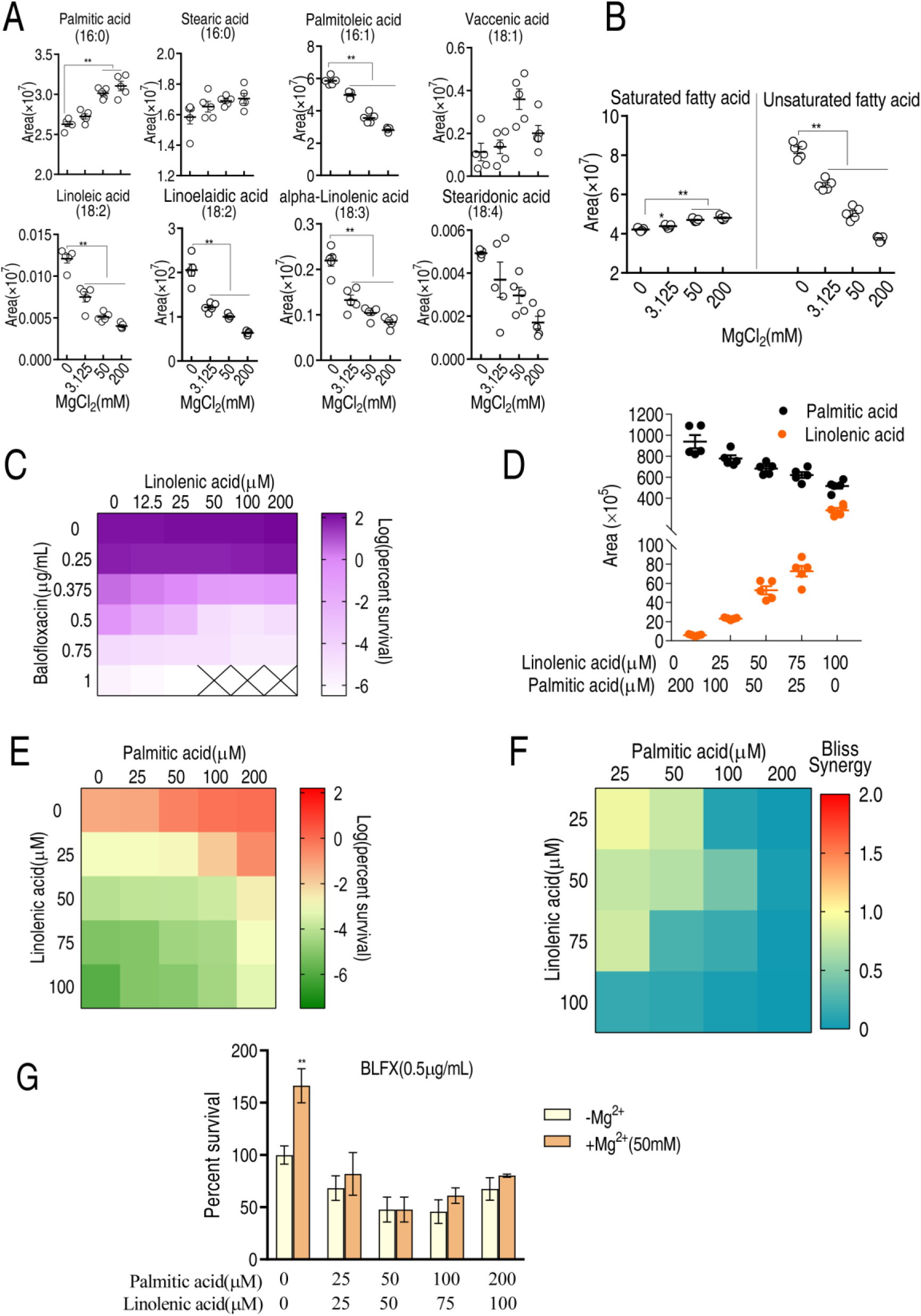
LC-MS targeted 16-carbon and 18-carbon fatty acids and the role of palmitic acids and linolenic acids in BLFX resistance. **A.** Scatter plots of 16-carbon and 18-carbon fatty acids, detected by LC-MS. The area indicates the area of the peak of the metabolite in total ion chromatography using GC-MS. **B.** Scatter plots of total saturated fatty acids and unsaturated fatty acids with 16-carbon and 18-carbon (**A**). **C.** Synergy analysis for BLFX with linolenic acid for ATCC 33787. Synergy is represented using a color scale or an isobologram, which compares the dose needed for 50% inhibition of synergistic agents (while) and non-synergistic (i.e., additive) agents (purple). **D.** LC-MS for the abundance of intracellular linolenic acid and palmitic acid of ATCC33787 in synergy with the indicated exogenous linolenic acid and palmitic acid. **E.** Synergy analysis of linolenic acid and palmitic acid in BLFX-mediated killing to ATCC33787. Synergy is represented using a color scale or an isobologram, which compares the dose needed for 50% inhibition of synergistic agents (blue) and non-synergistic (i.e., additive) agents (red). **F.** Bliss analysis **(E)**. **G**. Percent survival of ATCC33787 in the presence of linolenic acid, palmitic acid, and BLFX with or without 50 mM MgCl_2_.

Direct exposure to palmitic acid also inhibited balofloxacin-mediated killing (**Fig 2F**) while linolenic acid promoted balofloxacin-mediated killing in a dose-dependent manner (**Fig 4C**). Increasing exogenous palmitic acid and linolenic acid increased intracellular palmitic acid and linolenic acid levels, respectively (**Fig 4D**). When cells were co-treated with palmitic acid, linolenic acid, and balofloxacin, the two fatty acids appeared to antagonize each other, as demonstrated by the Bliss model (**Fig 4E** and **4F**). Furthermore, magnesium had a minimal effect on the antagonistic effect of palmitic acid, linolenic acid, and balofloxacin (**Fig 4G**), suggesting that this mineral functions through lipid metabolism. These results indicate that saturated and unsaturated fatty acids have an antagonistic effect on balofloxacin resistance.

### Mg^2+^ promotes phospholipid biosynthesis

Fatty acids are biosynthetic precursors of lysophosphatidic acid (LPA) and phosphatidic acid (PA), two key cell membrane components (Zhang and Rock, 2008a). Thus, LC-MS was used to assess the effect of Mg^2+^ on membrane lipid composition (**Suppl. Fig 7A**). Higher Mg^2+^ levels increased the percentage of lipids from 61 to 67%, and saturated fatty acids from 24 to 26%, but decreased the percentage of unsaturated fatty acids from 15 to 8% (**Suppl. Fig 7B**). The abundance of 11, 32, and 53 lipids was increased in 3.125, 50, and 200 mM MgCl_2_-treated bacteria, respectively, while the abundance of 26, 52, and 107 lipids was decreased in 3.125, 50, and 200 mM MgCl_2_-treated bacteria, respectively (**Suppl. Fig 7C**). Saturated fatty acids and lipids increased and unsaturated fatty acids decreased in an Mg^2+^ dose-dependent manner (**Suppl. Fig 7D**). A total of 52 lipids were quantified, including 17 high-abundance lipids (**Suppl. Fig 7E**). Phosphatidylethanolamine (PE) and phosphatidylglycerol (PG) had the first and second highest abundance (**Suppl. Fig 7E**) and PE levels declined while PG levels increased with increasing Mg^2+^ (**Fig 5A** and **5B**). A similar effect was observed for the unsaturated derivatives of PE and PG (**Fig 5C**). Principal component analysis (PCA) showed that component t [1] differentiated 0 and 3.125 mM Mg^2+^ from 50 and 200 mM Mg^2+^, while component t [2] separated 50 mM Mg^2+^ from 0 and 200 mM Mg^2+^ and variants in 3.125 mM of Mg^2+^ treatment (**Fig 5D**). S-plot analysis showed that behenic acid, PG[16:1(9Z)/16:1(9Z)], PG[18:2 (9Z, 12Z)/16:0], PA [14:0/22:6 (4Z, 7Z, 10Z, 13Z, 16Z, 19Z)], and lysoPE(16:0/0:0) were upregulated while linoelaidic acid, palmitoleic acid, PE-NMe(24:0/24:0), PE[16:0/14:1(9Z)], and lysoPA(i-12:0/0:0) were downregulated (**Fig 5E**). Of these, the abundance of PG[16:1(9Z)/16:1(9Z)], PG[18:2(9Z,12Z)/16:0], and behenic acid increased, while the levels of linoelaidic acid, palmitoleic acid, PE[16:0/14:1(9Z)], and lysoPA decreased with increasing Mg^2+^ levels (**Fig 5F**). Thus, altered phospholipid abundance may play a role in the effect of Mg^2+^ on antibiotic resistance.

**Figure 5.**
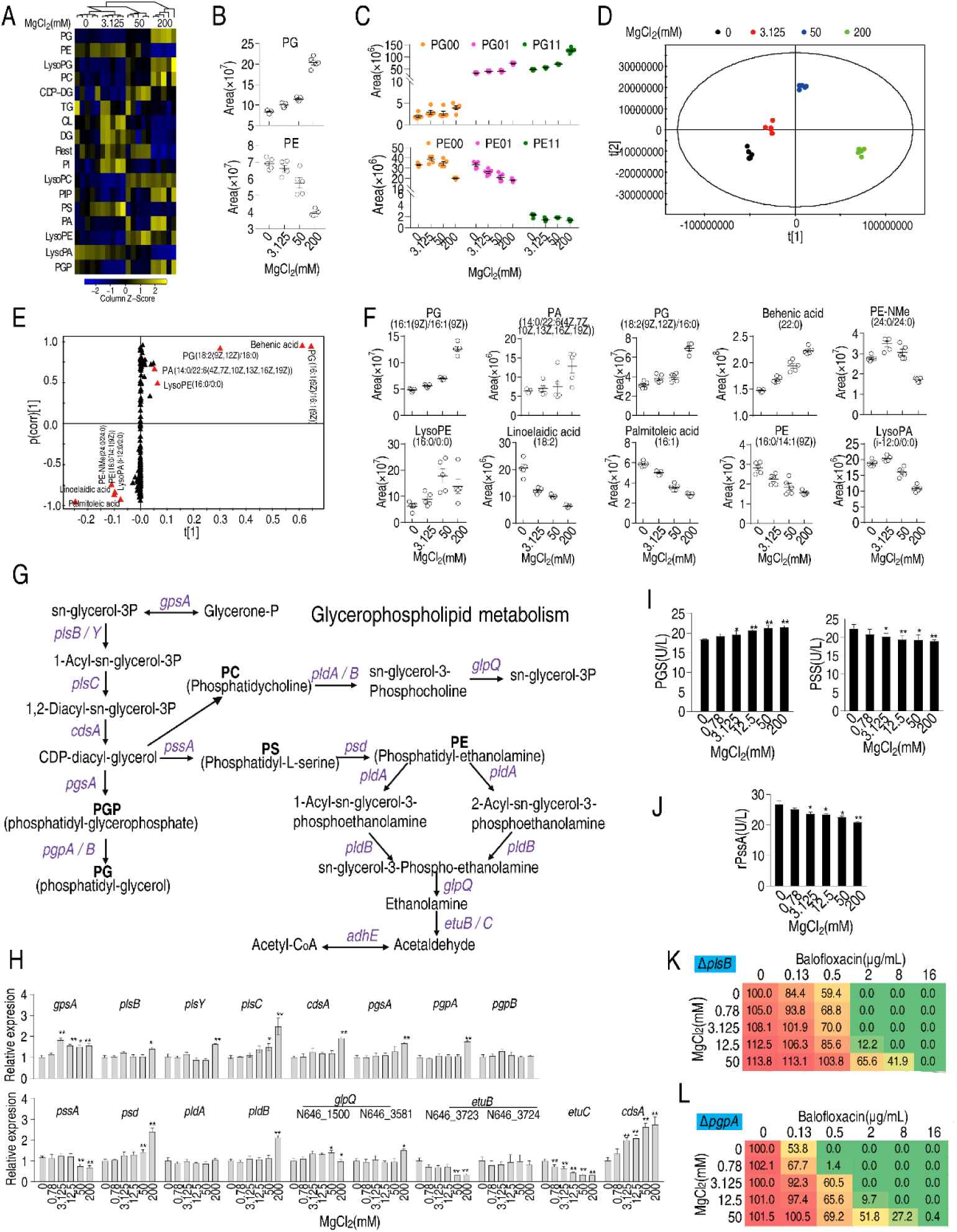
Effect of Mg2+ on phospholipid biosynthesis. **A.** Heatmap showing changes in differential lipid levels at the indicated concentration of MgCl_2_. **B.** Abundance of ATCC33787 phosphatidylglycerol (PG) and phosphatidylethanolamine (PE) at the indicated concentrations of MgCl_2_. **C.** Scatter plots of PG and PE at different saturation levels in the presence of the indicated MgCl_2_ levels. **D.** PCA analysis of different concentrations of MgCl_2_-induced phospholipids metabolomes. Each dot represents a technical replicate of samples in the plot. **E.** S-plot generated from OPLS-DA. Predictive component p [1] and correlation *p* (corr) [1] differentiated 0 and 3.125 mM MgCl_2_ from 50 and 200 mM MgCl_2_. **F.** Scatter plot of biomarkers in data (**E**). **G.** Diagram showing glycerophospholipid metabolism. **H.** qRT-PCR of the expression of genes encoding glycerophospholipid metabolism in the absence of or at the indicated concentrations of MgCl_2_. **I.** PGS and PSS levels in the absence or presence of the indicated concentrations of MgCl_2_. **J.** Activity of recombinant PSS in the absence or presence of the indicated concentrations of MgCl_2_. **K** and **L** Synergy analysis for MgCl_2_ with BLFX for Δ*plsB* (K) and Δ*pgpA* (L). Synergy is represented using a color scale or an isobologram, which compares the dose needed for 50% inhibition for synergistic agents (blue) and non-synergistic (i.e., additive) agents (red).

PE and PG are the two end products of phospholipid metabolism (**Fig 5G**). While expression of most genes in the phospholipid biosynthetic pathway, including *psd, pldB,* and *glpQ*, increased in the presence of 200 mM MgCl_2_, the expression of *pssA* and *etuB*, *etuC*, encoding the first enzyme and the last enzymes in the pathway, respectively, remained the same. In addition, the expression of *gpsA,* which encodes GpsA and transforms sn-glycerol-3P to glycerone-P, increased independent of Mg^2+^ concentration (**Fig 5H**). Phosphatidylglycerol phosphate synthase (PGS), encoded by *pgsA,* and phosphatidylserine synthase (PSS), encoded by *pssA,* are critical enzymes for the biosynthesis of PG and PE, respectively. PGS and PSS levels were elevated and reduced, respectively, with increasing Mg^2+^ (**Fig 5I**). The effect of Mg^2+^ on the activity of PSS was confirmed using recombinant PSS (**Fig 5J**). Deletion of *plsB* and *pgpA*, the first and last genes involved in PG biosynthesis, respectively, lowered cell viability in the presence of balofloxacin (note: Δ*psd* could not be obtained) (**Fig 5K** and **5L**). These results indicate that phospholipids may play a role in the effect of Mg^2+^ on balofloxacin resistance; more specifically, PE may inhibit and PG may promote resistance to balofloxacin.

### Mg^2+^ regulates membrane polarization, permeability and fluidity to confer balofloxacin resistance

Mg^2+^ remodels membrane composition, which may impair membrane potential and polarization, critical to membrane permeability and uptake of antibiotics (Lee *et al*., 2019; Peng *et al*., 2015). The voltage-sensitive dye, DiBAC4(3) showed that 12.5–200 mM MgCl_2_ promoted membrane depolarization in a dose-dependent manner (**Fig 6A**). Meanwhile, MgCl_2_ had a dose-dependent (**Fig 6B**) and time-dependent (**Fig 6C**) effect on proton motive force (PMF). These results suggest that Mg^2+^-dependent changes in membrane depolarization may influence antibiotic resistance.

**Figure 6.**
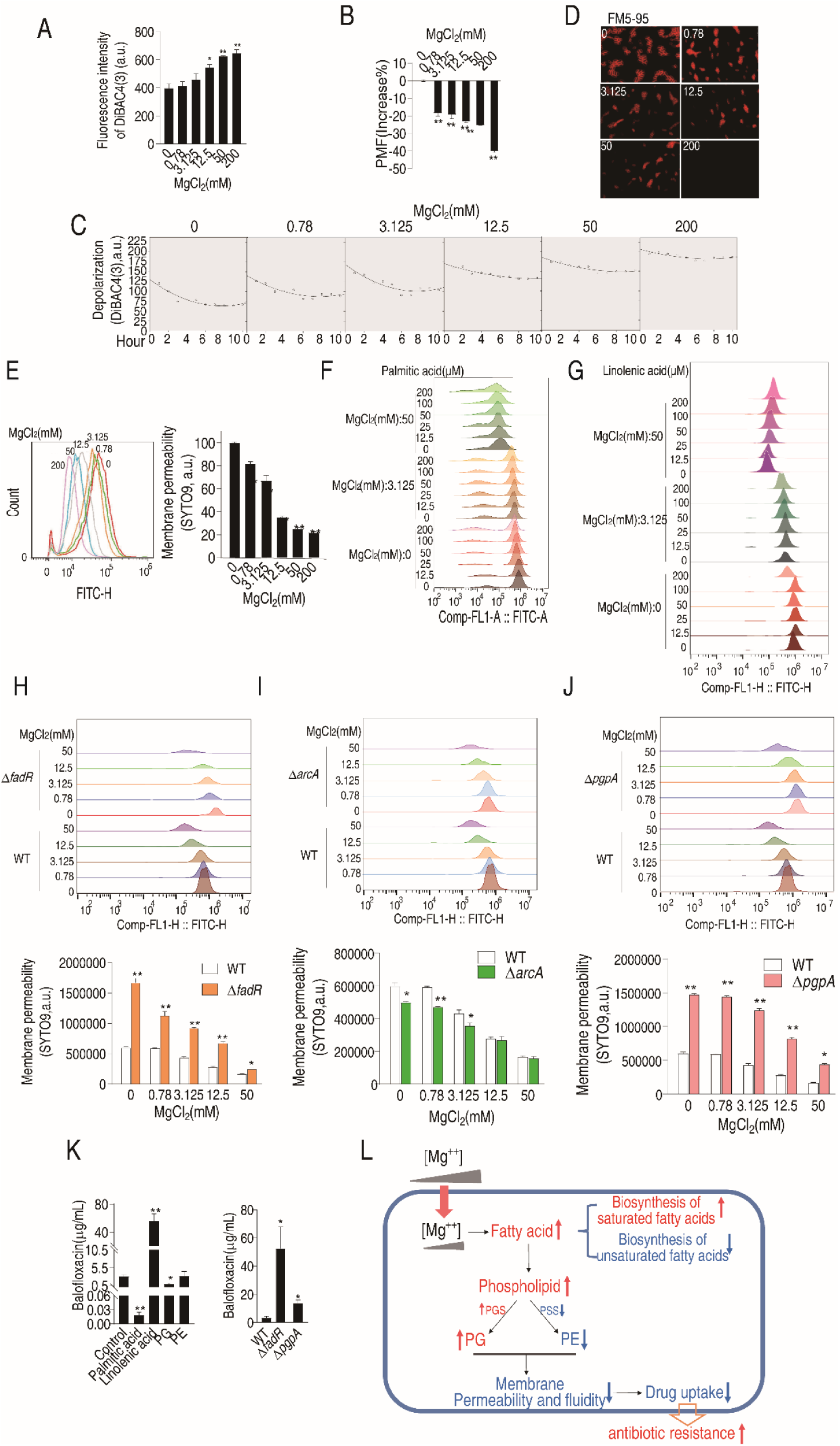
Mg^2+^ regulates membrane polarization, permeability, and fluidity to confer balofloxacin resistance. **A** and **B**. Depolarization **(A)** and PMF **(B)** of ATCC33787 in the absence of or at indicated concentrations of MgCl_2_. **C** and **D**. Dynamic depolarization (**D**) Membrane fluidity of ATCC33787 in the absence or presence of indicated concentrations of MgCl_2_, as shown by fluorescence microscopy. **E.** Membrane permeability of ATCC33787 in the absence or presence of the indicated concentrations of MgCl_2_. **F** and **G**. Membrane permeability of ATCC33787 cultured in palmitic acid **(F)** or linolenic acid **(G)** at the indicated concentrations of MgCl_2_. **H** and **I**. Membrane permeability of Δ*fadR* **(H)** and Δ*arcA* **(I)** in the absence or presence of the indicated concentrations of MgCl_2_. **J.** Membrane permeability of Δ*pgpA* in the absence or presence of the indicated concentrations of MgCl_2_. **K.** Intracellular BLFX of ATCC33787 in the presence of palmitic acid, linolenic acid, PG, and PE (left panel) or Δ*fadR* and Δ*pgpA* mutants (right panel). **L.** Diagram of mechanisms of Mg^2+^-mediated resistance to BLFX.

The efficacy of antibiotics is strongly influenced by bacterial membrane permeability and fluidity (Saeloh *et al*., 2018; Zhao *et al*., 2021). Thus, fluorescence microscopy was used to visualize these functions after incubating *V. alginolyticus* cells with FM5-95 and various concentrations of MgCl_2_. FM5-95 staining decreased with increasing concentrations of Mg^2+^, and no staining was observed in the presence of 200 mM Mg^2+^ (**Fig 6D**). SYTO9, a green fluorescent dye that binds to nucleic acid, enters and stains bacteria cells when there is an increase in membrane permeability (Lehtinen *et al*., 2004; McGoverin *et al*., 2020). Staining decreased with increasing MgCl_2_, indicating that bacterial membrane permeability declined in an Mg^2+^ dose-dependent manner (**Fig 6E**). These results indicate that exogenous Mg^2+^ reduces bacterial membrane permeability.

Exogenous palmitic acid also shifted the fluorescence signal peaks to the left in an MgCl_2_-dependent manner while palmitic acid only slightly shifted the peaks (**Fig 7F**). In contrast, exogenous linolenic acid shifted the peak to the right in a dose-dependent manner at 50 mM MgCl_2_ (**Fig 6G**). These results indicate that exogenous palmitic acid and linolenic acid decrease or increase membrane permeability, respectively. Membrane permeability was increased in Δ*fadR* cells than in WT cells and was reduced by Mg^2+^, remaining higher than the control (**Fig 6H**). However, Δ*arcA* cells had lower membrane permeability at lower MgCl_2_ concentrations (**Fig 6I**). Δ*pgpA* cells exhibited higher membrane permeability than control cells at all MgCl_2_ concentrations (**Fig 6J**). These data are consistent with the viability of the mutants described above (**Figs 3L** and **5L**).

Relative membrane permeability appeared to correlate with relative intracellular balofloxacin and antibiotic efficacy. Exogenous palmitic acid and linolenic acid reduced and promoted the uptake of balofloxacin at 159- and 18-fold, respectively (**Fig 6K**). Loss of *fadR* and *pgpA* increased intracellular balofloxacin (**Fig 6K**). These data suggest that bacterial membrane permeability is a critical factor required for the efficacy and uptake of balofloxacin, in the presence or absence of exogenous MgCl_2_.

## Discussion

This study explored the effect of Mg^2+^ on the phenotypic antibiotic resistance of *V. alginolyticus*. Exogenous Mg^2+^ was found to promote saturated fatty acid biosynthesis and inhibit unsaturated fatty acid biosynthesis. Mg^2+^ was also shown to upregulate PGS activity and PG abundance while downregulating PSS activity and PE abundance, decreasing membrane permeability and antibiotic uptake (**Fig 6L**) and enabling phenotypic resistance. These findings are consistent with the known impact of exogenous Mg^2+^ on bacterial survival in the presence of balofloxacin and other antibiotics, and the association between membrane permeability and drug efficacy. This study further showed that exogenous fatty acids influence membrane permeability and bacterial survival in the presence of balofloxacin.

Mg^2+^ is the most abundant divalent cation in cells (Pohland and Schneider, 2019; Pontes et al., 2015) playing many essential roles, including stabilizing macromolecular complexes and membranes, binding cytoplasmic nucleic acids and nucleotides, interacting with phospholipid head groups and cell surface molecules, and acting as an essential cofactor in many enzymatic reactions (Groisman *et al*., 2013). The present study showed for the first time that exogenous Mg^2+^ influences palmitic acid and linolenic acid levels and upregulates PGS while downregulating PSS. These findings extend our current understanding of the biological functions of Mg^2+^.

Recent findings suggest a link between fatty acid biosynthesis and the antibiotic resistance of *E. tarda* and *V. alginolyticus* (Su *et al*., 2021). The current study found that Mg^2+^ had opposing effects on the abundance of saturated and unsaturated fatty acids, stimulating saturated fatty acid biosynthesis at moderately high levels, while inhibiting unsaturated fatty acids biosynthesis at ≥3 mM. Thus, the ratio of saturated fatty acids to unsaturated fatty acids should help in predicting antibiotic resistance.

Prior studies of Gram-positive and Gram-negative bacteria (Kumariya *et al*., 2015; Said *et al*., 1987) found that 10–20 mM Mg^2+^ disrupts *Staphylococcus aureus* membranes and kills stationary-phase *S. aureus* cells but does not influence the survival of *Escherichia coli* and *Bacillus subtilis* (Xie and Yang, 2016). Low concentrations of Mg^2+^ (≤10 mM) induce PmrAB-dependent modification of lipid A in wild-type *E. coli* (Herrera *et al*., 2010). In two mundticin KS-resistant *Enterococcus faecium* mutants, a putative zwitterionic amino-containing phospholipid increased significantly, while phosphatidylglycerol and cardiolipin levels declined (Sakayori *et al*., 2003). However, the impact of Mg^2+^ on antibiotic resistance by regulating PGS and PSS activity was not reported. More importantly, exogenous MgCl_2_ was shown to target these enzymes in opposing ways to increase the PE/PG ratio, a previously unknown mechanism of Mg^2+^-mediated antibiotic resistance.

Prior studies show that in *Stenotrophomonas maltophilia,* PhoP/Q is activated by low magnesium levels and PhoP/Q inhibition or inactivation stimulates β-lactam antibiotic uptake (Huang *et al*., 2021). The effects of the outer membrane permeabilizers, polymyxin B nonapeptide and EDTA, are completely abolished by 3 mM Mg^2+^ (Kwon and Lu, 2006). In response to Mg^2+^-limited growth, enteric Gram-negative bacteria show higher lipid A acylation, which alters membrane permeability and reduces the uptake of cationic antimicrobial peptides (Guo *et al*., 1998). Mg^2+^ reduces the high sensitivity of rough *Salmonella typhimurium, S. minnesota*, and *Escherichia coli* 08 (which includes defects in the lipopolysaccharide carbohydrate core) mutants to several antibiotics (Stan-Lotter *et al*., 1979). Mg^2+^ (1 mM) inhibited aminoglycoside-mediated outer membrane permeabilization in *Pseudomonas aeruginosa* (Hancock *et al*., 1981). However, mechanisms underlying the impact of Mg^2+^ on antibiotic resistance have remained largely unknown. The current study revealed that Mg^2+^ plays a critical role in regulating the membrane permeability required for antibiotic resistance by modulating PE and PG during phospholipid metabolism.

Aquaculture is an environmental gateway to the development and globalization of antimicrobial resistance due to the excessive use of these drugs to prevent and treat bacterial contaminants (Cabello *et al*., 2020; Millanao *et al*., 2018). The current study found that Mg^2+^ increased antibiotic resistance, providing a reasonable explanation of why antibiotics are being excessively used in aquaculture and a possible solution to Mg^2+^-induced antibiotic resistance by balancing phospholipid metabolism. This finding should promote an overall reduction in antibacterial use that will improve environment and food safety.

In summary, Mg^2+^ modulated the phenotypic resistance of *V. alginolyticus* to balofloxacin and potentially other quinolones and other classes of antibiotics by altering membrane permeability and thus reducing antibiotic uptake. These findings inform the development of methods to improve the therapeutic effect of antibiotics on magnesium-induced phenotypic resistance.

## Materials and methods

### Bacterial strains and culture

*V. alginolyticus* ATCC33787 and *V. parahaemolyticus* 01 are from our laboratory collection (Jiang et al., 2022; Kou et al., 2022). They were cultured in 0.5% yeast broth (HuanKai Microbial, Guangdong, China) (pH 7.0) with 3% NaCl at 30 °C overnight. The cultures were diluted 1:50 (v/v) and grown in fresh 0.5% yeast broth supplemented with desired concentrations of MgCl_2_ at 30 °C until OD600 = 0.6. Bacteria were harvested by centrifugation at 8,000 rpm for 3 min and resuspended in corresponding concentrations of MgCl_2_ to 0.6 of OD600.

### Measurement of the minimum inhibitory concentration (MIC) by microtitre-dilution-method

MIC using microtitre-dilution-method was performed as previously described (Zhang et al., 2019). In brief, the overnight bacterial cultures in 3% NaCl were diluted at 1:100 (v/v) in fresh 0.5% yeast broth supplemented with desired concentrations of MgCl_2_, cultured at 30 °C and collected when the cells arrived at an OD600 of 0.6 in medium without MgCl_2_. Then, 1 X 10^5^ CFU cells were dispensed into each well of a 96-well microtiter polystyrene tray after which a series of 2-fold dilutions of antibiotic was added. Following incubation at 30 °C for 16 h, the MIC was defined as the lowest antibiotic concentration that inhibited visible growth. Three biological repeats were carried out.

### Measurement of MIC by Oxford method

Determination of MIC by Oxford cup was performed as previously described (Liu et al., 2015). Bacterial cells cultured overnight in yeast medium were diluted at 1:100, shaken at 30 ^0^C and 200 rpm until the OD_600_ nm reached at 0.6. Aliquot of 100 μL cells were spread on 0.5% yeast solid medium containing 3% NaCl and 0, 0.625, 1.25, 2.5, 5, 10, 20, or 40 mM of MgCl_2_. Oxford cups were placed on the solid medium, and various amounts of balofloxacin (0, 0.39, 0.78, 1.56, 3.125, 6.25, and 12.5 μg) were added. After culturing at 30 °C for 12 h, diameter of the inhibition zone was measured and the inhibition area was calculated.

### Measurement of intracellular balofloxacin

Measurement of intracellular balofloxacin was performed by LC-MS analysis and plate-counting assay. For LC-MS analysis, ATCC33787 cultured in the desired concentrations of MgCl_2_ were collected and adjusted to OD 0.6. Aliquot of 50 mL bacterial cells were added into a 250 mL Erlenmeyer flask and then 300 μL of 10 mg/mL balofloxacin (final concentration is 60 μg/mL). After being cultured for 6 h at 30 °C, these bacterial cells were collected and adjusted to OD 1.0 with 3% NaCl containing corresponding concentrations of MgCl_2._ Aliquot of 30 mL bacteria were collected and washed three times using mobile phase (acetonitrile: double distilled water containing 0.1mol/l formic acid = 35:65) for LC-MS detection. The bacterial cells were added 1 mL mobile phase, crushed in ice water bath for 10 min (crush for 2S, pause for 3S at 35% power). Following by centrifugation, supernatants were collected, filtered, and then measured by using liquid chromatography for balofloxacin. Different concentrations of balofloxacin were used for a standard curve. For plate-counting assay, ATCC33787 cells cultured overnight were diluted at 1:100, shaken at 30 ^0^C and 200 rpm until the OD_600_ nm reached at 0.6. The cells were diluted at 1:100 and spread on LB agar to preparing ATCC33787 plates. Meanwhile, ATCC33787 cells were cultured in medium with desired MgCl_2_ and at 30 °C and 200 rpm for 6 h. These cells were collected, washed and sonicated. The sonicated solution was added to Oxford cups with the ATCC33787 plates. After culturing at 30 °C for 12 h, diameter of the inhibition zone was measured. A gradient of balofloxacin was used as a standard curve for drug quantification.

### Measurement of intracellular magnesium

Measurements of intracellular Mg^2+^ concentration were carried out as previously described with a modification (Yang et al., 2018). In brief, *V. alginolyticus* cultured in 0%, 0.78%, 3.125%, 12.5%, 50%, and 200% MgCl_2_ were collected, washed, and resuspended in the same concentrations of MgCl_2_ to an OD600 of 1.0. Aliquots of 20 mL of bacterial suspensions were centrifuged and washed by ddH_2_O once. The resulting cells were weighted, which was designated wet weight. These samples were freeze-dried overnight, which was designated dry weight. Intracellular water volume was calculated using the following formula: W × (1-W/D-0.23) as described previously (Unemoto et al., 1973). Two hundred microliters of concentrated nitric acid was added to the dried cells and then heated in 75 °C for 20 min. After diluting the samples 20-fold, they were analyzed for Mg^2+^ concentration by inductively coupled plasma mass spectrometry (ICP-MS) (iCAP 6500, Thermo Fisher). The intracellular Mg^2+^ concentration was calculated according to the volume of intracellular water.

### Western blotting

Western blotting was carried out as described previously (Yao et al., 2019). Bacterial protein samples were prepared by ultrasound treatment, resolved on a 12% SDS-PAGE and transferred to nitrocellulose membranes (GE Healthcare Life Sciences). The membranes were incubated with 1:100 of the primary mouse antibodies, followed by goat anti-mouse secondary antibodies conjugated with horseradish peroxidase. Band intensities were detected by using a chemiluminescence imaging analysis system, Tanon-5200.

### Metabolomics analysis

Metabolomics analysis was performed by GC-MS as described previously (Yang et al., 2018). Briefly, ATCC33787 were cultured in 0.5% yeast broth with desired MgCl_2_. Equivalent numbers of cells were quenched with 60% (v/v) cold methanol (Sigma) and then centrifuged at 8,000 rpm at 4 °C for 5 min. One milliliter of cold methanol was used to extract metabolites. To do this, the samples were sonicated for 5 min at a 10-Wpower setting using the Ultrasonic Processor (JY92-IIDN, Scientz, China), followed by centrifugation at 12,000 rpm in 4 °C for 10 min. Supernatants were collected and 10 µL ribitol (0.1 mg per mL, Sigma-Aldrich, USA) was added into each sample as an internal quantitative standard. The supernatants were concentrated for metabolite derivatization and then used for GC-MS analysis. Every experiment was repeated by five biological replicates. GC-MS detection and spectral processing for GC-MS were carried out using the Agilent 7890A GC equipped with an Agilent 5975C VL MSD detector (Agilent Technologies, USA) as described previously (Unemoto et al., 1973; Yang et al., 2018). Statistical difference was obtained by Kruskal Wallis test and Manne Whitney test using SPSS 13.0 and a *p* value _<_ 0.01 was considered signi_fi_cant. Hierarchical clustering was completed in the R platform (https://cran.r-project.org/) with the function “heatmap. 2” of “gplots library”. Z score analysis was used to scale each metabolite. Multivariate statistical analysis included orthogonal partial least square discriminant analysis (OPLS-DA) implemented with SIMCA 12.0 (Umetrics, Umeå, Sweden). Control scaling was selected prior to fitting. All variables were mean centered and scaled to pareto variance of each variable. Rank-sum test and a permutation test were used to identify differentially expressed metabolites. OPLS-DA was used to reduce the high dimension of the data set. Differential metabolites to their respective biochemical pathways were outlined in the MetaboAnalyst 3.0 (http://www.metaboanalyst.ca/). Pathways were enriched by raw *p* value < 0.05.

### qRT-PCR

Quantitative real time polymerase chain reaction (qRT-PCR) was carried out as described previously (Yang et al., 2020). Total RNA was extracted from *V. alginolyticus* using TRIZOL regent (Invitrogen Life Technologies) according to the protocol. RNA Electrophoresis was carried out in 1% (w/v) agarose gels to identify quality of the extracted RNA. By using a PrimeScriptTM RT reagent Kit with gDNA eraser (Takara, Japan), reverse transcription-PCR was carried out on 1 μg of total RNA and primers are listed in **Table EV2.** qRT-PCR was performed in 384-well plates with a total volume of 10 μL and the reaction mixtures were run on a LightCycler 480 system (Roche, Germany). Data are shown as the relative mRNA expression compared to 0% MgCl_2_ test with the endogenous reference 16S rRNA gene.

### Antibiotic bactericidal assay

Antibiotic bactericidal assay was performed as described previously with a modification (Li et al., 2016; Zhao et al., 2021). The cultured bacteria of ATCC33787 and its mutant strains were transferred to fresh yeast medium at a dilution of 1:1000 and dispensed into test tubes and then the indicated concentrations of MgCl_2_ and balofloxacin were added. If desired, 2 mM 2-aminooxazole and 1 μg/mL triclosan were or a metabolite complemented. These mixtures were incubated at 30 °C and 200 rpm for 6 h. Cells were collected. To determine CFU per mL, 100-μL samples were 10-fold serially diluted with 900 μL M9 buffer and an aliquot of 5 μL of each dilution was spotted onto the LB agar plates and cultured at 30 °C for 8 h.

### Construction of gene-deleted mutants

Construction of gene-deleted mutants was carried out as described previously (Kuang et al., 2021). Primers were designed as shown in **Table EV3** using CE Design V1.03 software. To construct gene-deleted mutants, upstream and downstream 500-bp fragments were amplified from the genome using two pairs of primers (primers P1 and P2, primers P3 and P4), and then merged into a 1,000-bp fragment by overlap PCR using a pair of primers (primers P1 and P4). After the fragments were digested by *Xba*I, they were ligated into the pDS132 vector digested by the same enzymes, transformed into MC1061 competent cells. Conversion products were coated on LB plate containing 25 μg/mL chloramphenicol and cultured at 37 ℃ overnight. The plasmids from colony growing on the plate were identified by PCR using a pair of primers (primers P1 and P4) and sequenced. The sequenced plasmids were transformed into MFD λ pir competent cells as donor. MFD λpir and recipient bacterium ATCC 33787 were cultured to an optical density (OD) of 1.0 and then mixed at a ratio of 4:1. After centrifugation, the pellets were resuspended with 50 μL LB medium including DAP (100 μg/mL), dropped onto sterilized filter paper on LB medium including DAP (100 μg/mL), and cultured for 16-18 h at 37 °C.

All bacteria rinsed from the filter paper with LB medium were smeared onto the LB plate with chloramphenicol (25 μg/mL) and Ampicillin (100 μg/mL). After cultured for 16-18 h at 37 °C, bacteria were screened by LB plate with above two antibiotics. The bacteria were identified by plasmid PCR using a pair of primers P1 and P4 and sequencing and then were cultured and smeared onto the LB plates with 20% sucrose. The clones were cultured and smeared onto the LB plates with 20% sucrose again. The clones, which did not grow on the LB plates with chloramphenicol but grew on the LB plates with 20% sucrose, were identified by PCR using primer P7P8 (primer P7 is set at about 250 bp upstream of P1, and primer P8 is set at about 250 bp downstream of P4), P4P7, and P5P6 (for amplification of the target gene).

### LC-MS analysis for lipidomics

ATCC33787 were cultured in medium with desired concentrations of MgCl_2_, washed by 3% NaCl with the desired concentrations of MgCl_2_, and adjusted to OD 1.0. Aliquot of 30 mL the cultures was centrifuged each sample and bacterial cells were collected. Aliquot of 800 μL distilled water was added and treated for 5 min at 100 °C to inactivate phospholipase C. Protein concentration was determined with BCA kit. Sample solution with 3 mg protein was transferred into a 10 mL centrifuge tube, and supplemented with distilled water to 800 μL. Then all samples were processed as follows: 1 mL chloroform and 2 mL methanol (to make the volume ratio of chloroform: methanol: water = 1:2: 0.8) were added and cholic acid was as an internal standard; These mixtures were vortexed for 2 min and then 1 mL chloroform was add and vortex for 30s; 1 mL 10% NaCl solution was add, vortexed for 30s, and placed at room temperature overnight. After the solution was layered, a 2.5 mL syringe was used to suck the lowest layer (chloroform layer) to a new EP tube. Solution of the chloroform layer was evaporated in the rotary evaporator for 2 h and then 1 mL mobile phase (50% and 50%B. A, 25 mM ammonium acetate / methanol (30: 70); B, methanol) was added for analysis of lipids by LC-MS. Database (https://hmdb.ca) was used for lipid identification.

### Measurement of enzyme activity

Activity of pyruvate dehydrogenase (PDH), α-ketoglutarate dehydrogenase (KGDH) and succinate dehydrogenase (SDH) was carried out as previously described (Jiang et al., 2020). Cells cultured in medium were collected and washed three times with saline. The bacterial cells were suspended in Tris–HCl (pH 7.4) and disrupted by sonic oscillation for 6 min (200 W total power with 35% output, 2 s pulse, 3 s pause over ice). After centrifugation, supernatants were collected. The protein concentration of the supernatant was determined using Bradford assay (Beyotime, P0009). Then, 200 μg proteins were used for determination of pyruvate dehydrogenase (PDH), α-ketoglutarate dehydrogenase (KGDH) and succinate dehydrogenase (SDH) activity. Levels of PGS and PSS were quantified by ELISA kits according to manufacture’s instruction (Shanghai Fusheng Industrial Co., Ltd., China)

### Measurement of proton motive force

Measurement of the transmembrane voltage delta PSI (δPSI), which is the electrical component of the PMF, was performed as previously described (Cheng et al., 2018). Bacteria cultured in medium with desired concentrations of MgCl_2_ were collected at centrifugation and labeled by DiO2(3). Approximately 1 × 10^7^ CFU were added to a flow cytometry tube containing 1 mL buffer with 10 μM DiOC2(3) (Sigma) and incubated in the dark for 30 min at 30 ^0^C. Samples were assayed with BD FACSCalibur flow cytometer with a 488 nm excitation wavelength. Gates for bacterial populations were based on the control population by using forward versus side scatter and red versus green emission. Size and membrane potential determined the intensity of Red (488 nm excitation, 610 nm emission) fluorescence. The diverse ratios of red and green indicated fluorescence intensity values of the gated populations. Computational formula of membrane potential: 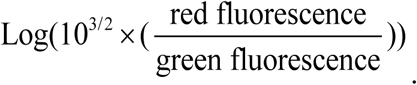.

### Measurement of depolarization

The membrane potential was estimated by measuring the fluorescence of the potential-dependent probe DiBAC4(3) (Saint-Ruf et al., 2016). DiBAC4(3) is a s voltage-sensitive probe that penetrates depolarized cells, binding intracellular proteins or membranes exhibiting enhanced fluorescence and red spectral shift. ATCC33787 were cultured in medium with desired concentrations of MgCl_2_, collected and then adjusted to OD 0.6. Aliquot of 100 μL bacterial cells were diluted to 1 mL, and 2 μL of 5 mM DiBAC4(3) was added. After incubated for 15 min at 30 ℃ without light and vibration, these samples were filtered and detected by BD FACSCalibur.

### Measurement of fluidity by fluorescence microscopy

Measurement of membrane fluidity is performed as previously described (Wen et al., 2022). Briefly, ATCC33787 were cultured in medium with indicated concentrations of MgCl_2_, collected and then adjusted to OD 0.6. Aliquot of 100 μL bacteria cells of each sample were diluted to 1 mL and 10 μL (10 mg/mL) FM5-95 (Thermo Fisher Scientific, USA) was added. FM5-95 is a lipophilic styryl dye that insert into the outer leaflet of bacterial membrane and become fluorescence. This dye preferentially bind to the microdomains with high membrane fluidity(Wen et al., 2022). After incubated for 20 min at 30 ℃ at vibration without light, the sample was centrifuged for 10 min at 12,000 rpm. The pellets were resuspended with 20 μL of 3% NaCI. Aliquot of 2 μL sample was dropped on the agarose slide, and take photos under the inverted fluorescence microscope.

### Measurement of membrane permeability

Measurement of membrane permeability was carried out as described previously (Su et al., 2021). ATCC33787 were cultured in medium with desired concentrations of MgCl_2_, collected and then adjusted to OD 0.6. Aliquot of 100 μL bacteria cells of each sample were diluted to 1 mL and 2 μL 10 mg/mL SYT09 was added. After incubated for 15 min at 30 ℃ at vibration without light, the mixtures were filtered and measured by flow cytometry (BD FACSCalibur, USA).

## Acknowledgments

This work was sponsored by the International Cooperation and Exchange program of National Natural Science Foundation of China (32273177) (to P.B.), Innovation Group Project of Southern Marine Science and Engineering Guangdong Laboratory (Zhuhai) (311020006) (to P.B.), Natural Science Foundation of Guangdong Province (2022A1515012079) (to L.H.), the Science and Technology Planning Project of Guangdong Province (2023B1212060028), and Fundamental Research Funds for the Central Universities, Sun Yat-sen University (24lgzy004).

## Author contributions

B. Peng conceived the idea, designed the experiments, and supervise the projects. H. Li, J. Yang, and S.F. Kuang conducted the experiments and interpreted the results.

## Supplementary Materials for

### Supplementary text

#### Magnesium and balofloxacin binding only partially contributes to phenotypic resistance

Prior studies suggest that the chelation of antibiotics by magnesium ions inhibits antibiotic uptake (Deitchman et al., 2018; Lunestad and Goksøyr, 1990). To investigate whether magnesium binds to balofloxacin, balofloxacin was pre-incubated with magnesium, and zone of inhibition (ZOI) analysis was conducted. Six different concentrations of balofloxacin were separately incubated with six different concentrations of MgCl_2_, and then spotted on filter paper so that a defined amount of balofloxacin could be used for ZOI. While lower concentrations of MgCl_2_, (0.78, 3.125, or 12.5 mM) did not alter the ZOI, higher concentrations, including 50 and 200 mM MgCl_2_, decreased the ZOI (**Suppl. Fig 2A**), suggesting that even high doses of magnesium had only a partial effect on balofloxacin through direct binding. For example, at 200 mM MgCl_2_ and 5 or 10 μg/mL balofloxacin, the balofloxacin ZOI was 53.2 and 70.3% of the ZOI at 0 mM MgCl2, suggesting that ≥50% of the antibiotics were still functional. Intracellular BLFX also decreased with increasing MgCl_2_ (**Suppl. Fig 2B**), while exogenous Mg^2+^ increased intracellular Mg^2+^ levels in a dose-dependent manner. For example, exogenous 50 and 200 mM MgCl_2_ increased intracellular Mg^2+^ levels to 1.21 and 1.31 mM, respectively (**Suppl. Fig 2C**). The relationship between TolC, an efflux pump that transports quinolones from bacterial cells, and Mg^2+^ was also assessed (Kobylka *et al*., 2020; Song *et al*., 2020). The expression of TolC/*tolC* was unaffected by Mg^2+^ (**Suppl. Fig 2D**). Magnesium is critical for LPS stability. LPS levels increased at 200 mM Mg^2+^ (**Suppl. Fig 2E**), however, the loss of *waaF, lpxA*, and *lpxC*, three key genes involved in LPS biosynthesis, did not influence balofloxacin sensitivity/resistance in the presence of Mg^2+^ (**Suppl. Fig 2F**). These findings suggest that magnesium-induced LPS biosynthesis does not contribute directly to BLFX resistance and demonstrate that Mg^2+^ influx is involved in balofloxacin resistance.

### Supplementary Tables

**Table 1.** Comparison in components in LBS and ASWT.

|  | **LBS** | **ASWT** |
| --- | --- | --- |
| Tryptone(%) | 1 | 1 |
| Yeast<br>extract(%) | 0.5 | 0.5 |
| NaCl(mM) | 171 | 210 |
| MgSO<sub>4</sub>(mM) | 0 | 35 |
| KCl(mM) | 0 | 7 |
| CaCl<sub>2</sub>(mM) | 0 | 7 |

**Table 2.**
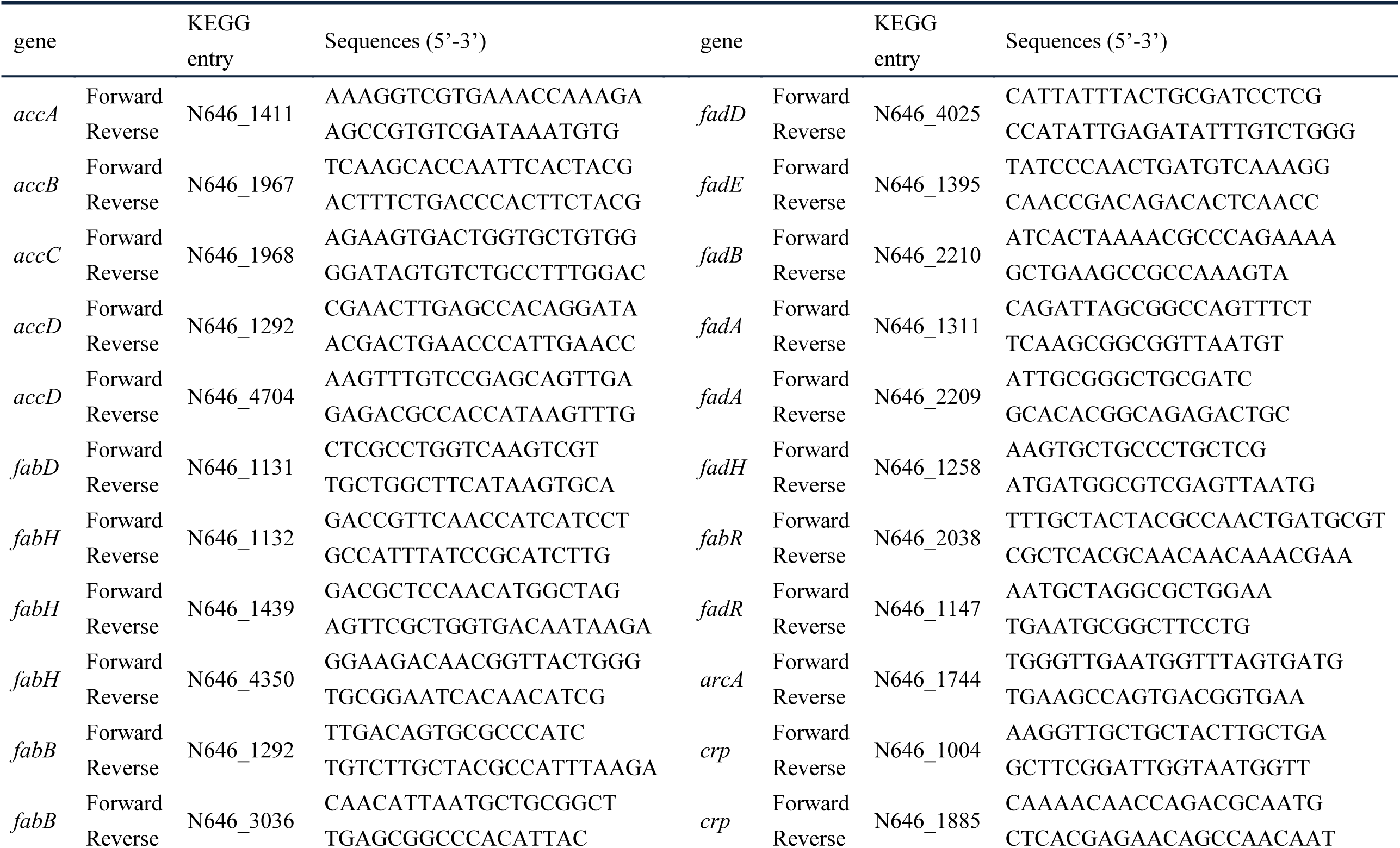
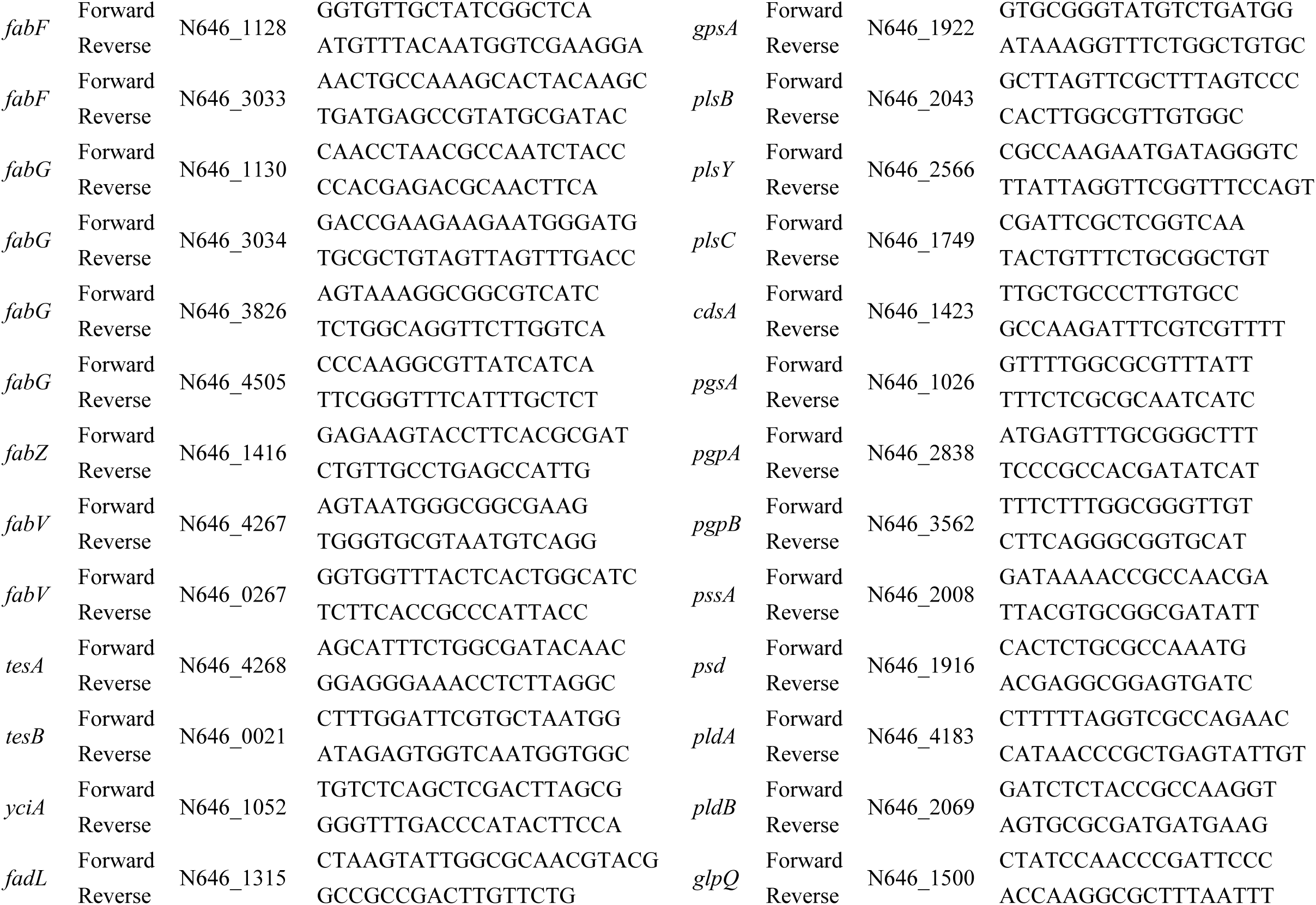
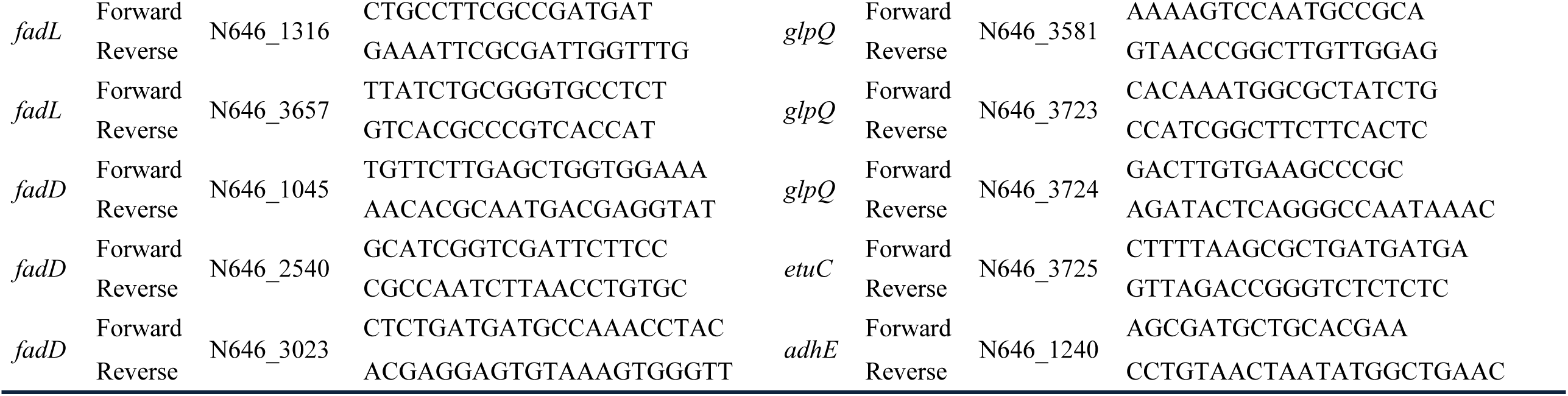
Primes used in the present study.

**Table 3.**
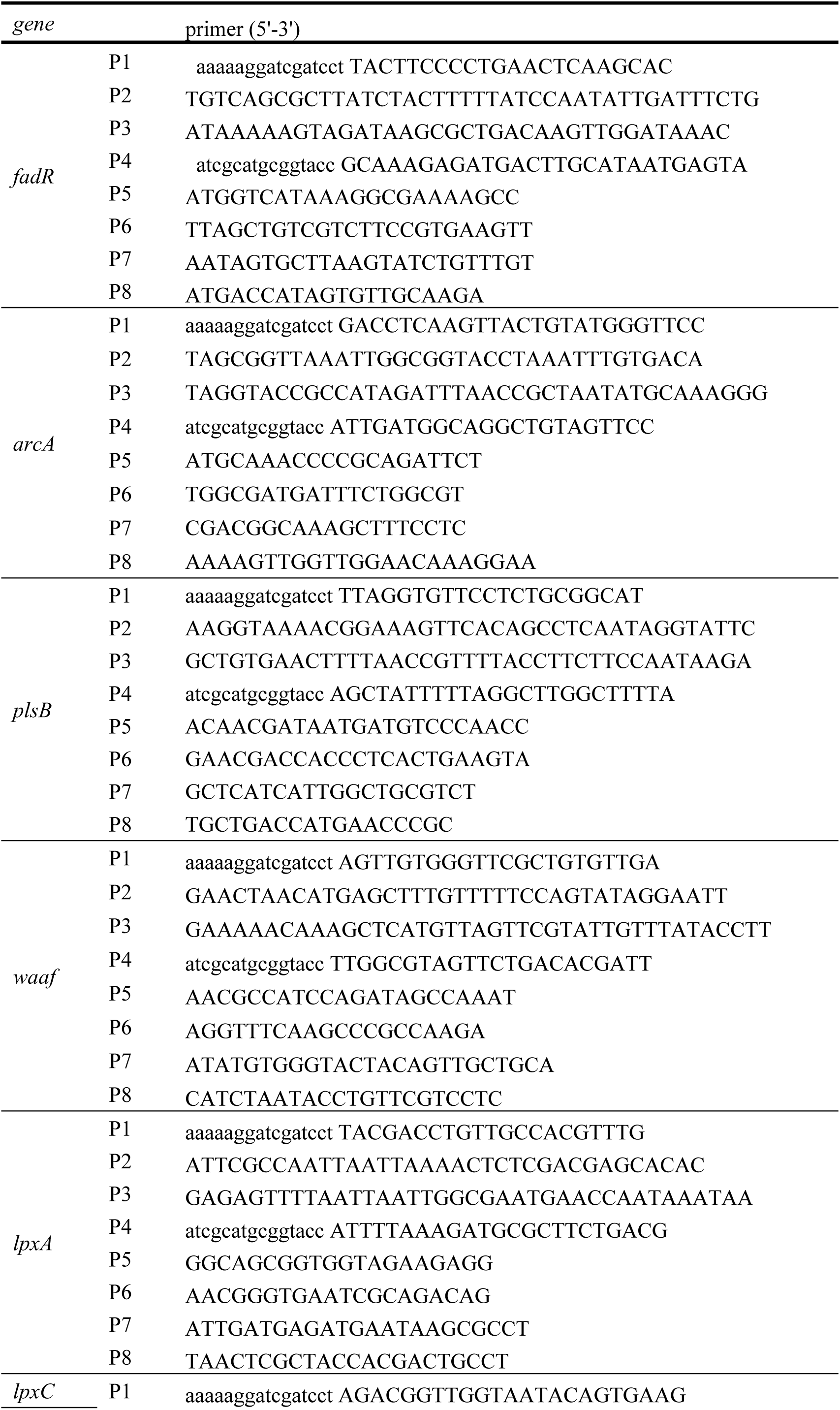
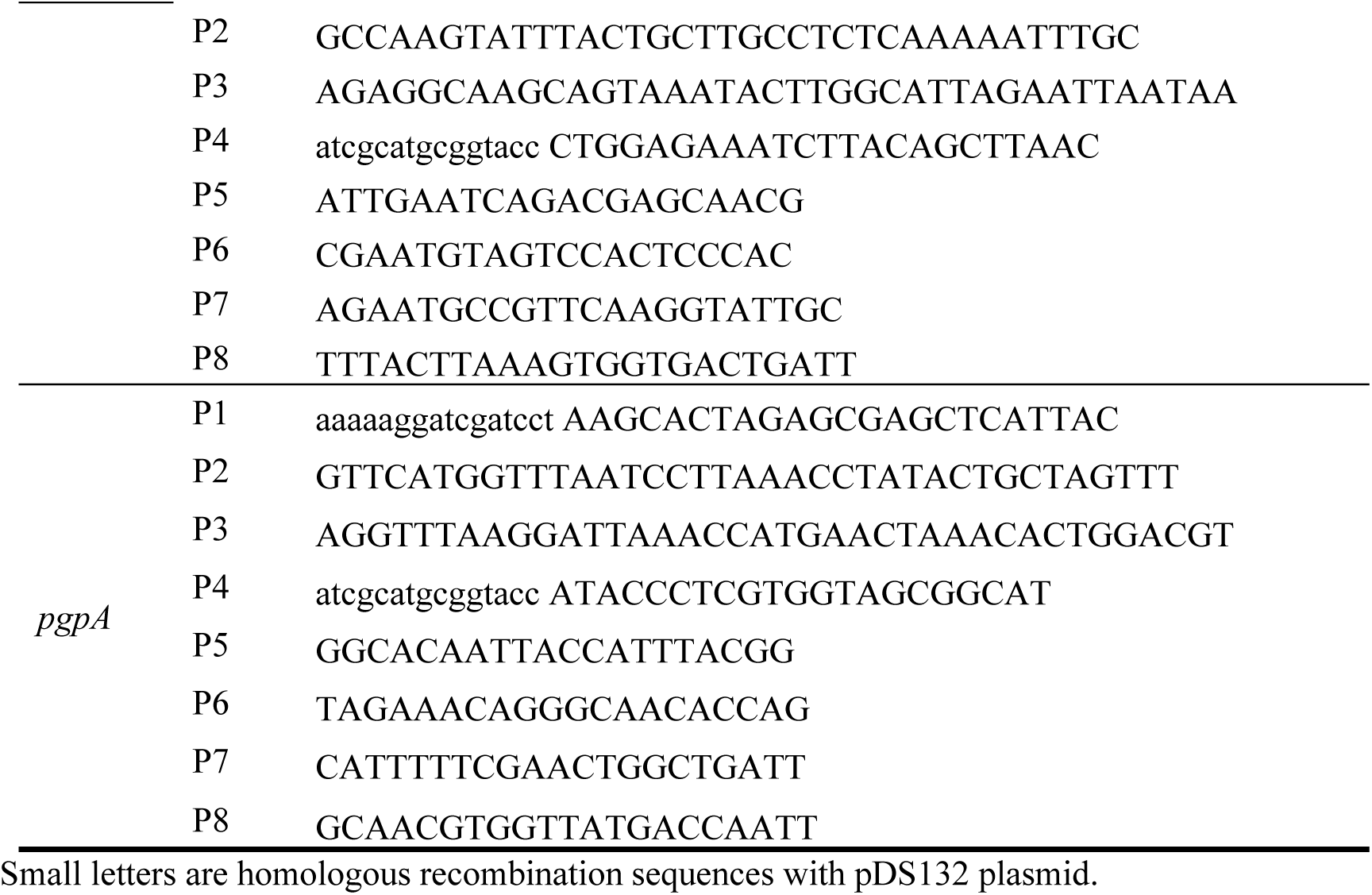
Primers used in the present study for construction of gene-deleted mutants.

### Supplementary Figures

**Figure 1.**
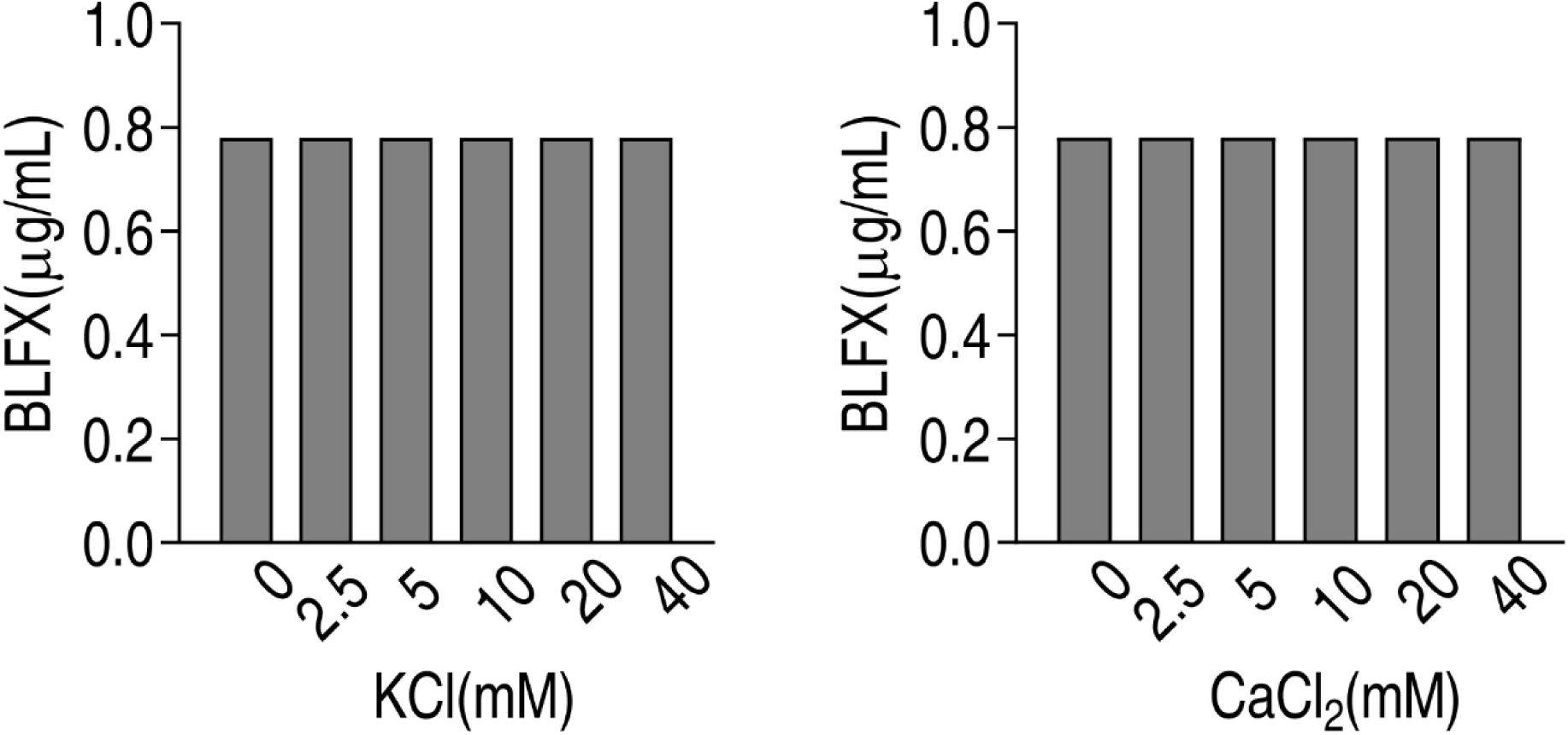
MIC of ATCC33787 to BLFX in the absence or presence of the indicated concentrations of KCl or CaCl_2_ in M9 media.

**Figure 2.**
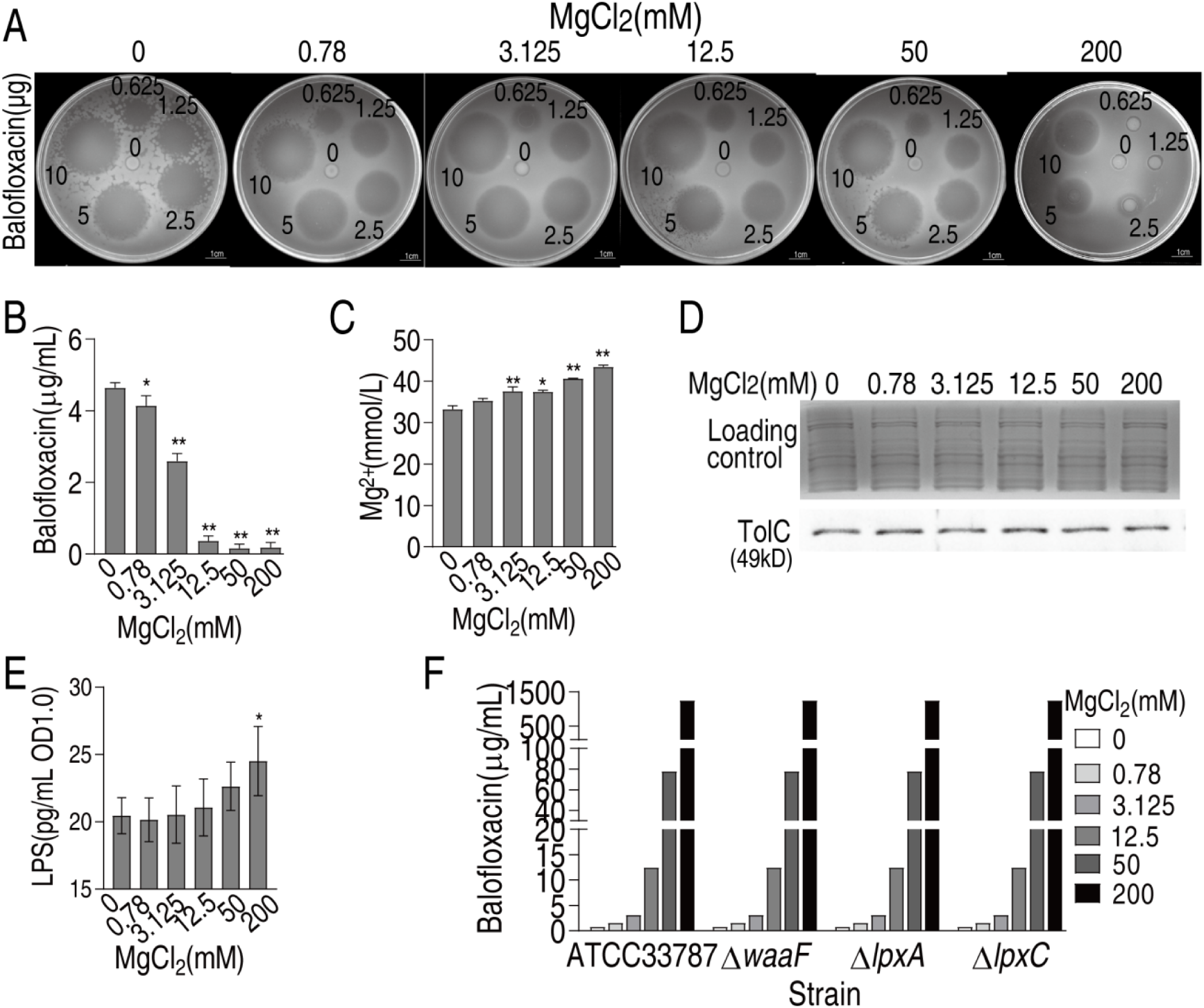
Mg^2+^ decreases balofloxacin uptake. A. Oxford cup test for effect of different concentrations of Mg^2+^ on antibacterial action of different dose of BLFX to ATCC33787. To do this, 0 mM, 0.78 mM, 3.125 mM, 12.5 mM, 50 mM, and 200 mM MgCl_2_ were individually mixed with 0 μg/mL, 12.5 μg/mL, 25 μg/mL, 50 μg/mL, 100 μg/mL or 200 μg/mL BLFX for 5 h. Then, 50 μL were added into Oxford cup for antibacterial efficiency, which contained 0 μg, 0.625 μg, 1.25 μg, 2.5 μg, 5 μg, or 10 μg BLFX, respectively. B. Intracellular BLFX of ATCC 33787 in ASWT with the indicated concentrations of MgCl_2_ and 60 μg/mL BLFX. C. Intracellular Mg^2+^ of ATCC 33787 in ASWT with the indicated concentrations of MgCl_2_. D. Western blot for abundance of TolC in the presence of MgCl_2_. Whole cell lysates resolved by SDS-PAGE gel was stained with Coomassie brilliant blue as loading control. E. LPS quantification at indicated concentrations of MgCl_2_. F. MIC of ATCC 33787 and its mutants Δ*waaF*, Δ*lpxA*, Δ*lpxC* in ASWT with the indicated concentrations of MgCl_2_, which is measured by microtitre-dilution-method.

**Figure 3.**
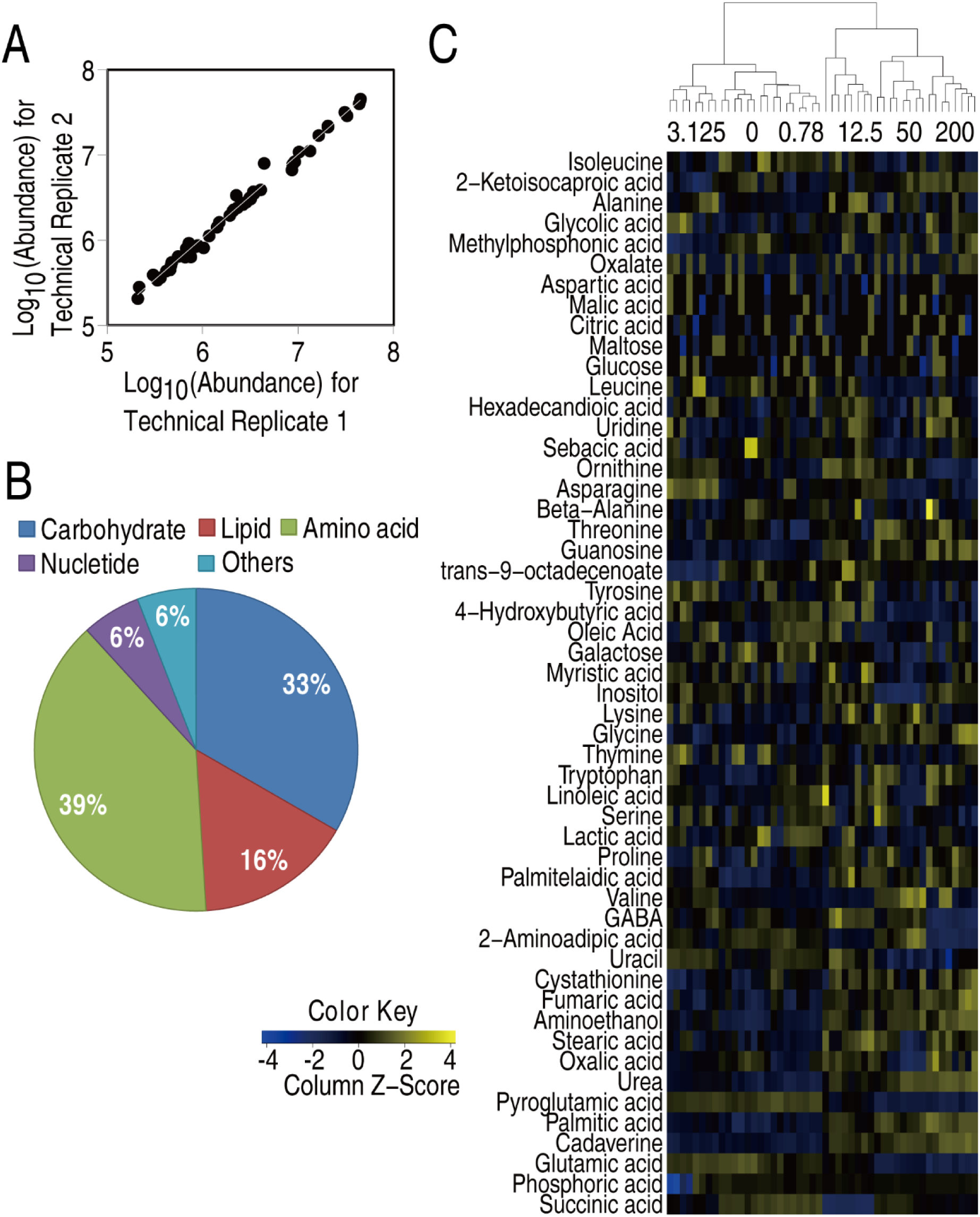
Metabolic profiles of *V. alginolyticus* in different concentrations of MgCl_2._ **A.** Reproducibility **B.** Percentage of metabolites in every category **C.** Heatmap of metabolites

**Figure 4.**
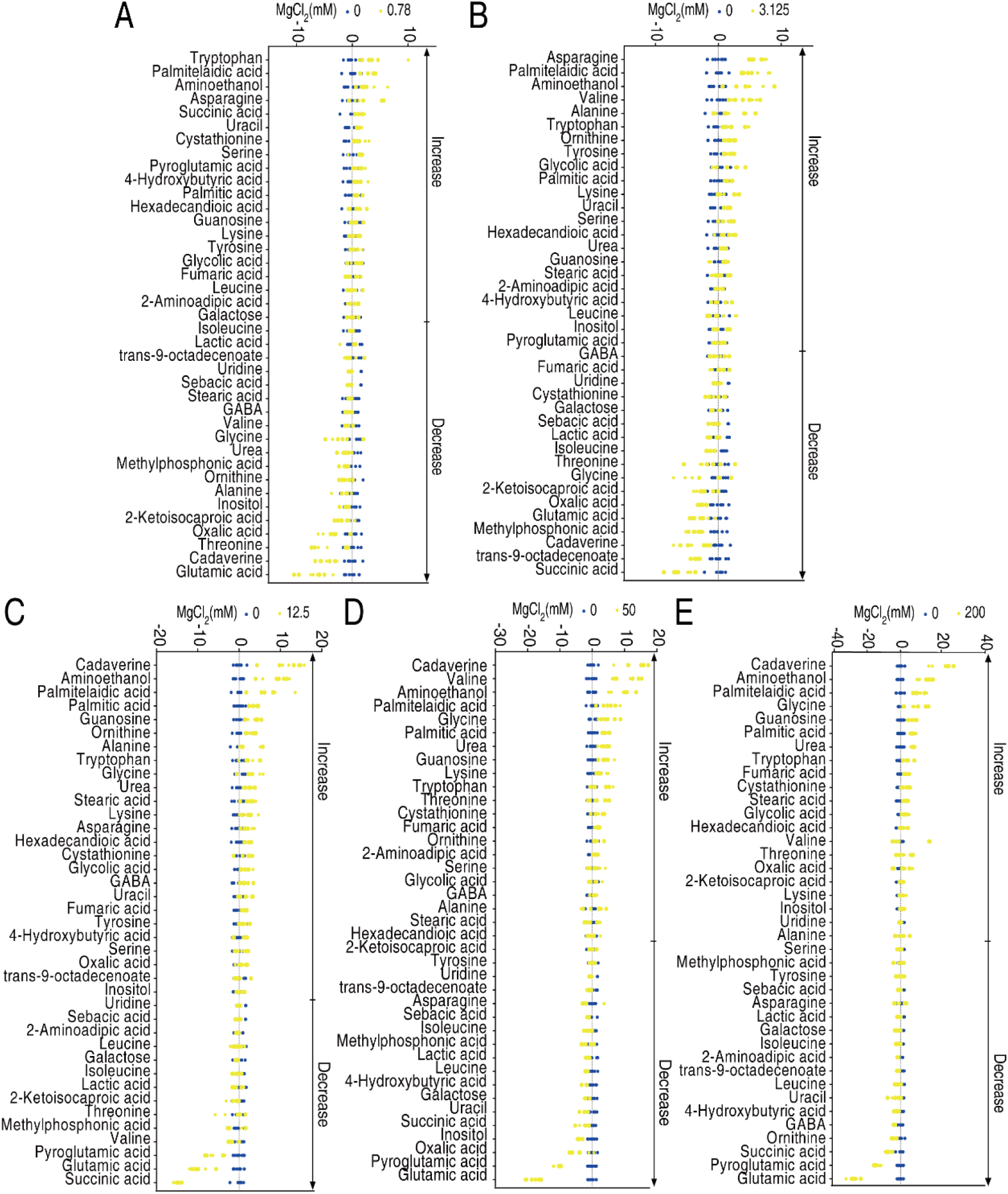
Heatmap and Z score plots of differential metabolites. **A.** Heatmap of differential metabolites **B-F**. Z score plots of differential metabolites.

**Figure 5.**
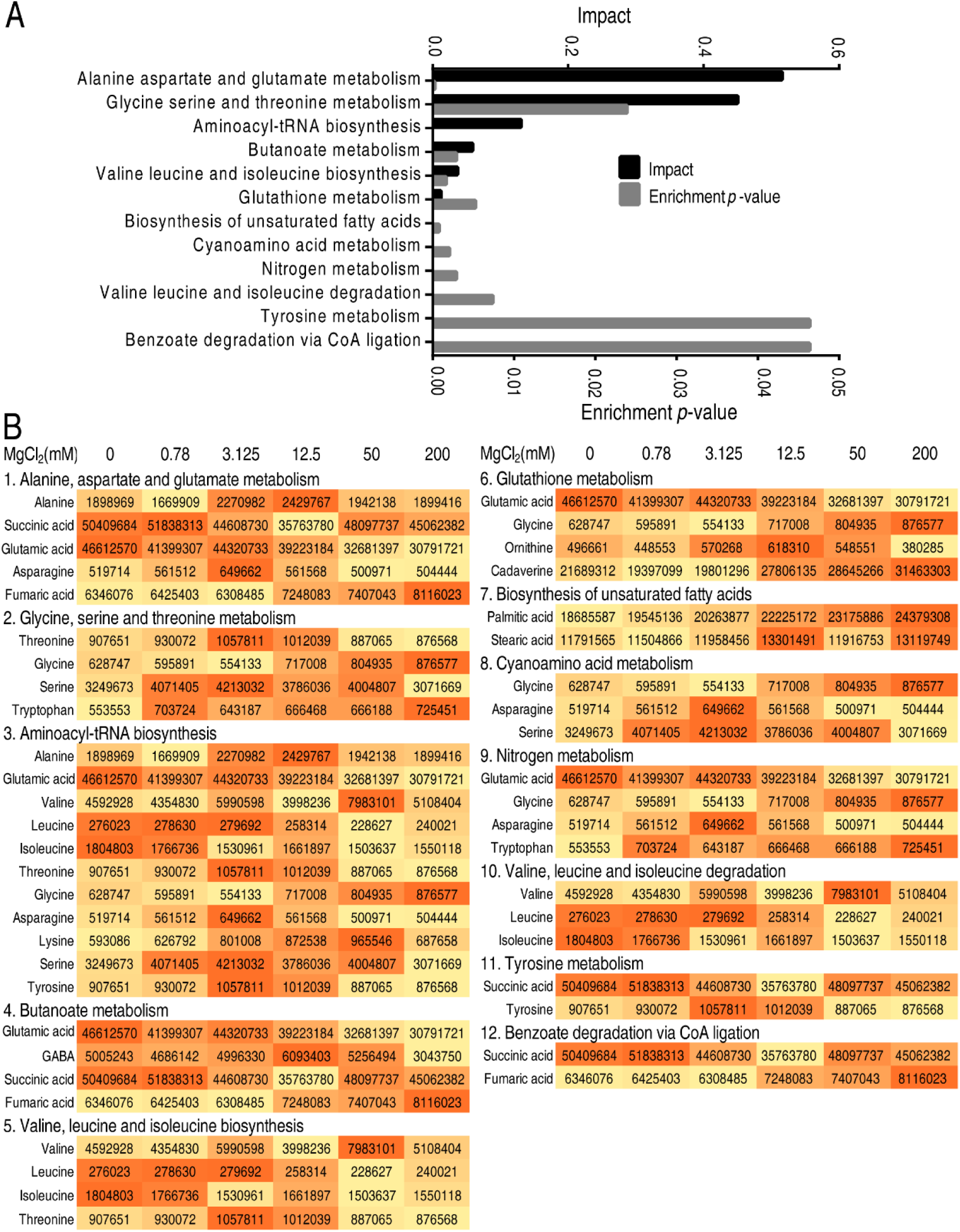
Pathway enrichment of differential metabolites. **A.** Pathway enrichment of differential metabolites **B.** Differential metabolites in enriched pathways

**Figure 6.**
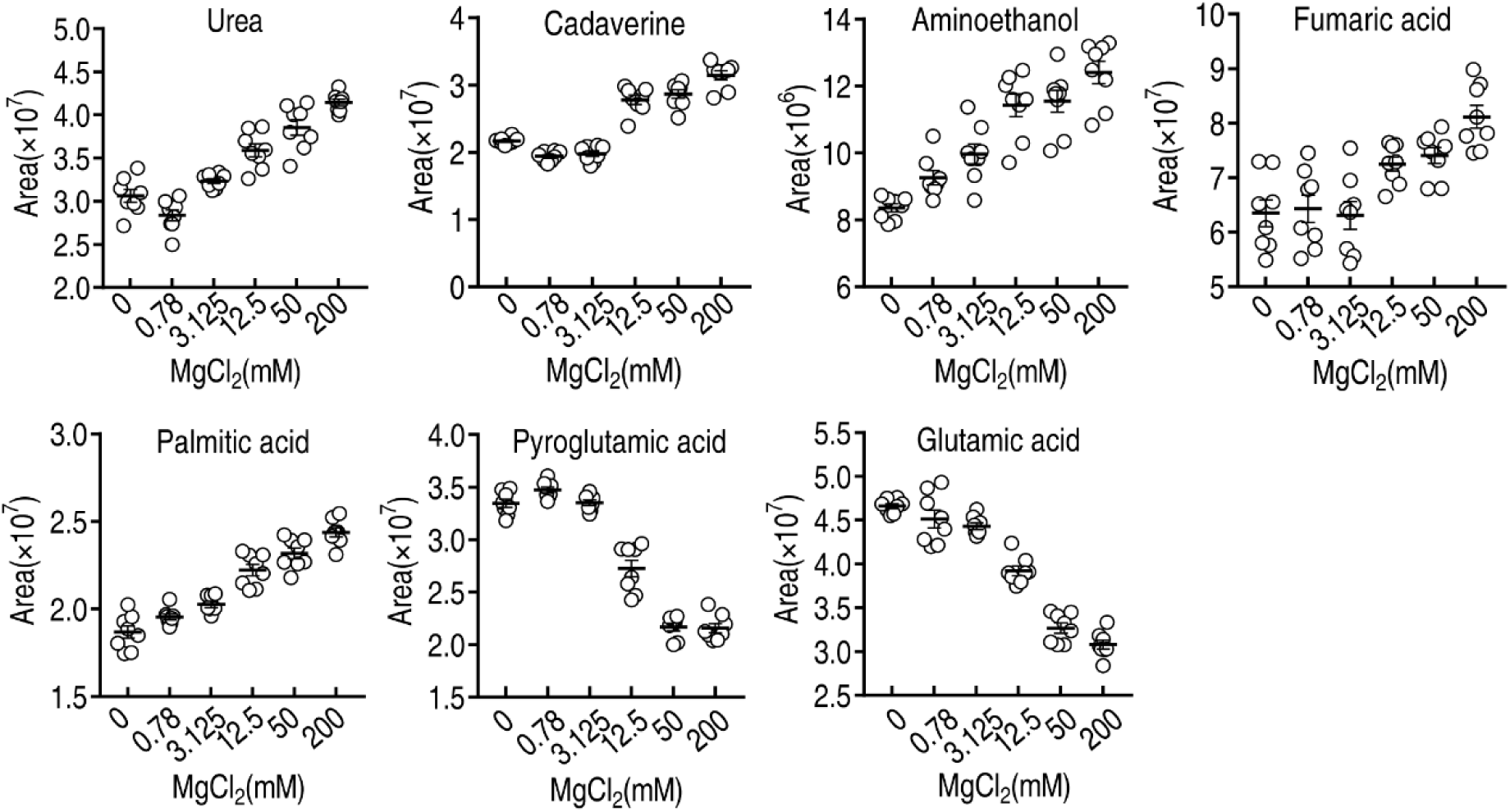
Scatter plots of differential metabolites identified by S-plot.

**Figure 7.**
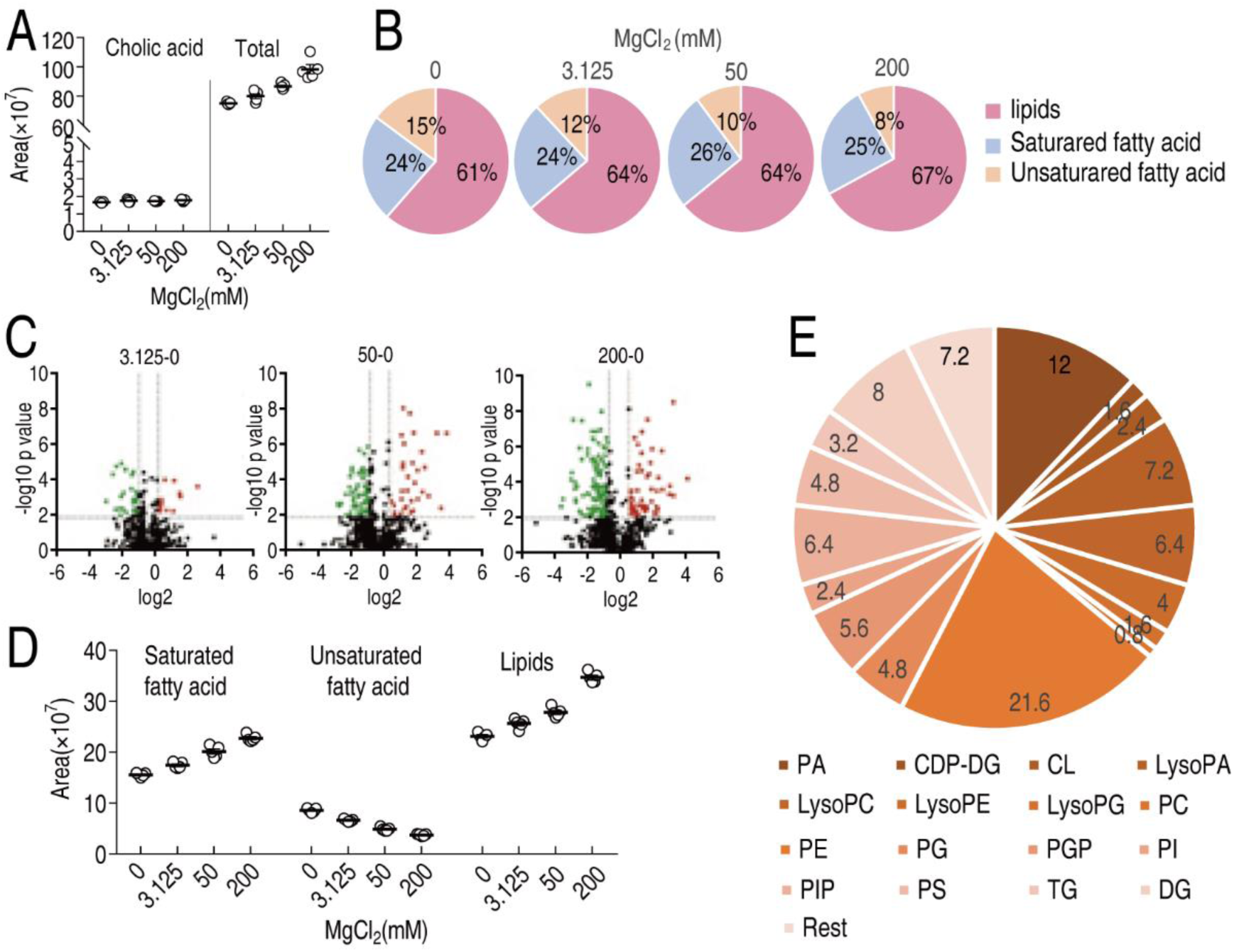
Lipidomes in the different concentrations of MgCl_2_. A. Area of fatty acids in the presence of indicated concentrations of MgCl_2_. B. Percentage of lipids, saturated fatty acid and unsaturated fatty acid in the presence of indicated concentration of MgCl_2_. C. Volcano plots of lipidomics of indicated concentration of MgCl_2_ as compared to non-treated control. D. Relative abundance of saturated fatty acids, unsaturated fatty acids and lipids in the presence of indicated concentrations of MgCl_2_. E. Relative percentage of indicated lipids.

